# A novel music-based real-time fMRI neurofeedback interface modulates interhemispheric connectivity and enhances mood

**DOI:** 10.1101/2025.05.06.652265

**Authors:** Alexandre Sayal, João Pereira, Bruno Direito, Teresa Sousa, Miguel Castelo-Branco

## Abstract

Music is a universal language that transcends cultures and is deeply rooted in human evolutionary history. Its creation and appreciation recruit the limbic and reward systems, leading to the evocation of emotions ranging from happiness and sadness to tenderness and grief. Here, we investigate the potential of music as an interventional tool in a novel neurofeedback connectivity-based experiment. This study proposes a musical interface for real-time fMRI neurofeedback that is adaptable to diverse experimental paradigms, namely the ones aiming at improving mood and other affective dimensions. Using a previously developed motor imagery connectivity-based approach, we evaluate its feasibility and efficacy by comparing the modulation of bilateral premotor cortex (PMC) activity during functional runs with real versus sham (random) feedback in 22 healthy adults. We also assess its performance against a visual feedback interface. The experiment involves a 50-minute MRI session, including anatomical scans, a PMC functional localizer run, and four neurofeedback runs (two with active feedback and two with sham feedback). Pre- and post-session questionnaires assess the neurobehavioral impact on mood, musical background (as a potential predictor of NF success), and subjective feedback experiences. During neurofeedback, participants perform motor imagery of finger-tapping, with feedback delivered as a dynamic, pre-validated chord progression that evolves or regresses based on the functional connectivity between left and right PMC. We found that our implementation of music-based feedback was successful, with participants managing to modulate their own connectivity using the proposed interface. The modulation performance was similar for active and sham NF runs, possibly due to the power of music to boost neuromodulation, but the network recruitment was stronger for active NF, including in the insula, putamen, and target ROIs. Behaviorally, we found a decrease in tension and an improvement in the overall mood of the participants after the session. When comparing our results to previous NF data with a visual interface, we found stronger brain activations, in particular in NF-relevant regions such as the insula and the putamen. This work shows that it is possible to directly modulate interhemispheric connectivity using a rt-fMRI musical interface with direct effects on mood and recruitment of saliency and learning networks.

## Introduction

Neurofeedback (NF) is an innovative brain-computer interface (BCI) technique, enabling individuals to modulate their brain activity through real-time feedback. With applications ranging from cognitive enhancement to clinical therapy, NF holds significant potential for improving mental health and well-being in psychiatric and neurological conditions (***Direito et al., 2021***; ***Sitaram et al., 2017***; ***Young et al., 2021***). However, advancing NF methodologies to optimize usability and effectiveness in real-world therapeutic settings remains a critical challenge for researchers (***Paret et al., 2019***).

The wide adoption of NF interventions is hindered by several challenges. A significant barrier is the high proportion of non-responders - individuals who fail to achieve consistent modulation of their brain activity - resulting in highly variable success rates across studies (***Kadosh and Staunton, 2019***). Another challenge lies in the methodological heterogeneity of NF research, including inconsistencies in defining appropriate control groups for different study objectives (***Sorger et al., 2019***) and the lack of standardized reporting for key success metrics (***Ros et al., 2020***). Addressing these issues is crucial for improving NF’s reliability and scalability in clinical and real-world settings.

Among the many factors influencing NF success, the design of the feedback interface - the medium through which neuronal information is conveyed to participants - plays a critical role (Figure 1). The type and presentation of feedback can strongly affect how participants engage with the task, influencing both the clarity of the signal and the subjective experience of control and immersion (***Direito et al., 2019***; ***Ninaus et al., 2013***; ***Stoeckel et al., 2014***). Optimizing the interface design, using neuroscience-oriented approaches, may therefore help reduce the proportion of non-responders by enhancing participant engagement and the effectiveness of the feedback loop. We raised the hypothesis that using music as a strong modulator of emotion and reward circuits, one might achieve a higher likelihood of neuromodulation success. In this study, we aimed to design and validate a real-time fMRI NF approach based on modulation of connectivity using music as the interface between participants and their own brain activity - a step towards impacting affective processing and mood.

**Figure 1.**
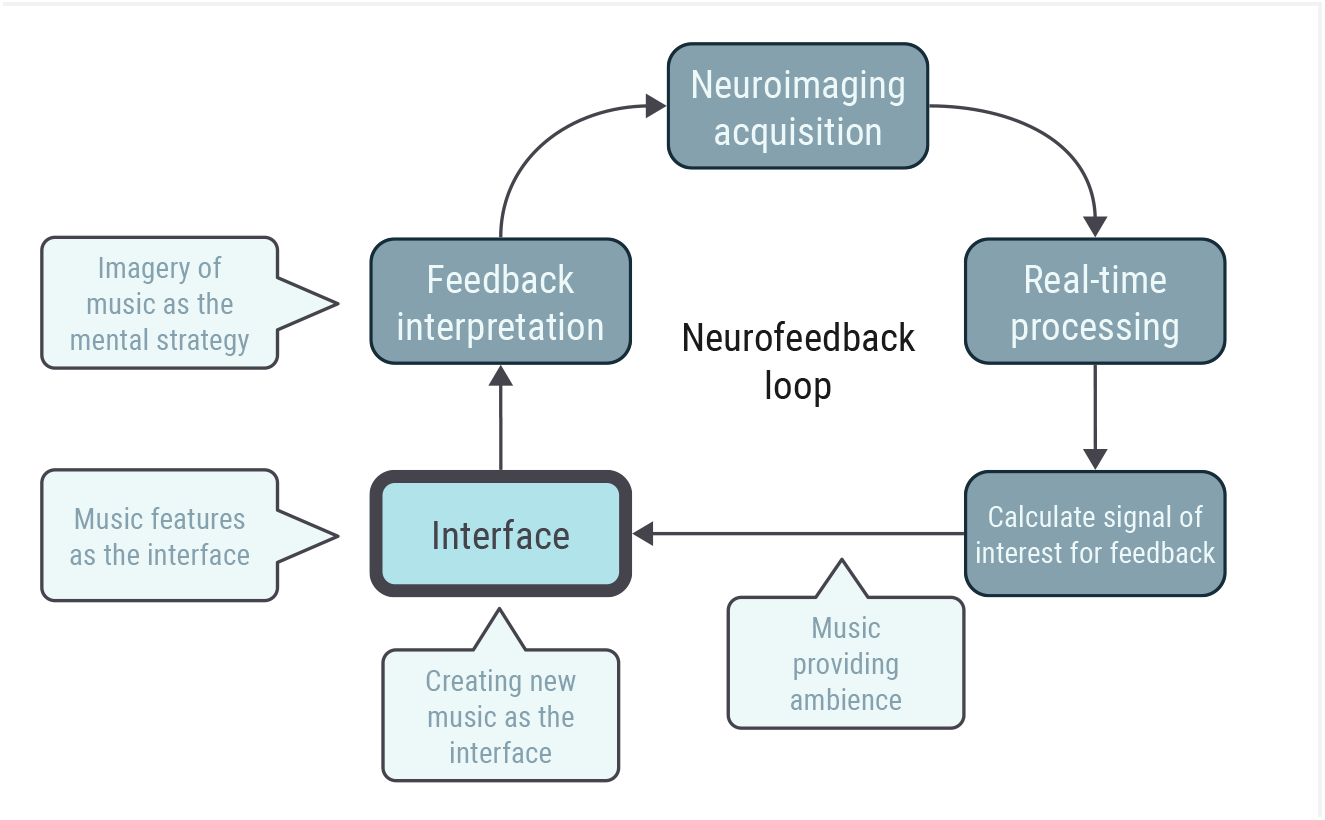
Illustration of the neurofeedback loop. The participant’s brain activity is acquired, analyzed, and transformed into feedback via an interface, which the participant uses to adjust their neural activity in real time. Music can play a role at various points in this process, from being the feedback to influencing participant engagement and interpretation. Here, we focus on the interface. Adapted from (***Sayal et al., 2025***).

Music is a universal phenomenon, deeply rooted in human biology, culture, and history, with the ability to evoke and regulate emotions across contexts (***Zatorre, 2024***). Listening to music consistently engages the brain’s limbic and reward systems, enabling it to elicit both basic emotions, such as happiness and sadness, and complex ones, like tenderness or grief. In fact, the sound encoding mechanisms recruited during music listening and interpretation are based on a large and hierarchical network of brain areas, including primary and secondary auditory cortices, anterior cingulate cortex, insula, hippocampus, somatosensory cortex, and basal ganglia (***Bonetti et al., 2024***; ***Ferreri et al., 2019***; ***Koelsch, 2020***; ***Koelsch et al., 2021***), regions tightly linked to the emotion circuits (***Barrett, 2017***).

As a tool for emotion regulation, music allows individuals to modulate their emotional states by either suppressing undesired feelings or inducing new ones (***Cook et al., 2019***). Moreover, extensive evidence links music listening and training to enhanced brain plasticity, highlighting its potential as an effective medium for neurofeedback interventions (***Blum et al., 2017***; ***Matziorinis et al., 2024***; ***Vik et al., 2018***).

Prior studies have explored the use of music in the context of NF approaches, and we recently conducted a systematic review to assess these effects (***Sayal et al., 2025***). We investigated the primary motivations for incorporating music, the methodological approaches employed, and the reported outcomes. Many studies emphasized music’s ability to engage neural circuits related to emotion and reward, but causal explanations of these effects were still lacking. Moreover, no consensus was found on the imaging or behavioral metrics used to define neurofeedback success when music was involved. Focusing on whole-brain neural correlates of music stimuli and their interactions with target brain networks and reward mechanisms when designing music-based neurofeedback interventions can help to address these gaps. Recent efforts to decode brain patterns associated with interpreting valence and arousal (***Daly, 2023***; ***Putkinen et al., 2021***; ***Sayal et al., 2024***) or complex emotions (***Koelsch et al., 2021***) in music add to our understanding of the underlying neural mechanisms, providing insights for the design of new music-based feedback approaches.

In this study, we developed a sham-controlled NF experiment with a novel interface based on musical chord progressions and implemented it in a real-time fMRI setup. Building on a previous NF experiment by our group, which established the feasibility of connectivity-based feedback during a motor imagery task (***Pereira et al., 2019***) using a visual thermometer interface, we created a novel setup to explore music as the feedback medium. Our primary goal was to assess the feasibility of using connectivity and music-based feedback for real-time fMRI NF interventions. To this end, we investigated the following research questions:

1. Can participants successfully modulate both the activity and interhemispheric correlation of the premotor cortices (PMC) using music-based feedback?
2. Does the music-based NF session positively affect participants’ mood?
3. Do participants accurately identify active NF runs (with contingent feedback) as more effective in terms of performance and feedback contingency compared to sham runs (with random feedback)?
4. Does contingent feedback elicit greater activation in reward- and learning-related brain regions compared to random feedback?
5. Aiming for a more immersive interface and a neuroscience-oriented approach, are the brain activation metrics stronger for the music feedback than for other approaches, such as the ones based on visual interfaces?

## Methods and Materials

### Study design overview

The study design is summarized in Figure 2. We followed the CRED-nf guidelines (***Ros et al., 2020***) to ensure transparent and rigorous reporting of our neurofeedback protocol; the completed checklist is provided in the Supplementary Materials.

**Figure 2.**
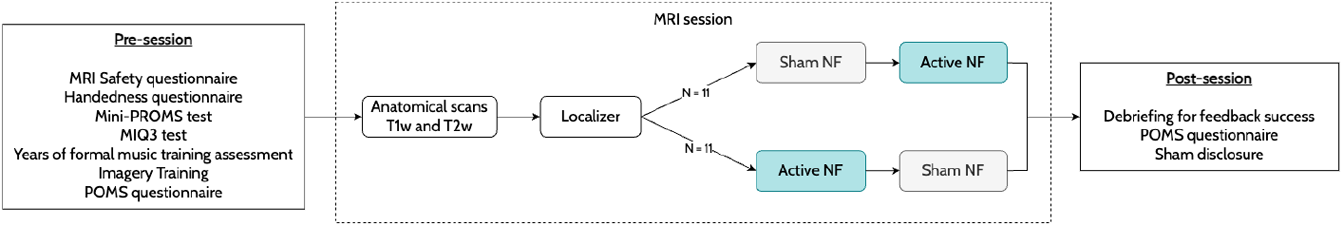
The experimental design includes a magnetic resonance imaging (MRI) session encompassing anatomical scans, a localizer run for the PMC, and four music-based neurofeedback runs (two with active feedback and two with sham). Participants completed pre- and post-session questionnaires to assess MRI safety, musical training, mood, and their subjective experiences regarding the feedback contingency.

After recruitment and before the MRI acquisition session, participants completed a series of questionnaires. The first questionnaire assessed each participant’s compatibility with the MRI environment to ensure safety and compliance. Next, we characterized the sample based on three parameters: musical training, handedness, and motor imagery ability. Participants completed the Mini-PROMS questionnaire (***Zentner and Strauss, 2017***), which assessed musical ability, and provided the number of years of formal music training. To assess handedness, we used a questionnaire (***Cohen, 2008***) based on (***Oldfield, 1971***). To evaluate motor imagery ability, participants completed the MIQ-3 questionnaire (***Mendes et al., 2016***; ***Williams et al., 2012***), which screens for visual and kinesthetic imagery capabilities.

Participants then watched a video explaining the neurofeedback experiment. This video detailed the number and type of sequences, the paradigm of the two tasks, and included a guided opportunity to practice motor imagery while listening to feedback music. To assess the impact of the NF session on mood, participants completed the POMS questionnaire (***Faro Viana et al., 2001***) both before and after the MRI session. The scoring was obtained for six subscales: tension, depression, hostility, fatigue, confusion, and vigor.

The MRI session lasted approximately 50 minutes and included the acquisition of anatomical scans, a functional localizer run, and four NF runs (two active and two sham). A crossover design was implemented: half of the participants received sham feedback during the first two runs, while the other half received sham feedback during the last two runs. The feedback shown during the sham NF runs was randomly generated.

After the MRI session, participants were asked to describe their experiences, including their perceived performance and feedback contingency during the runs. They were also asked to identify the best and worst runs regarding these parameters. Finally, the runs in which they received sham feedback were disclosed.

### Participants

A sample of 22 healthy participants (13 females, mean age 33 ± 6 years, range 24-42 years) was recruited for this study. The research plan adheres to the legal regulations of the General Data Protection Regulation (GDPR), national legislation, and the Declaration of Helsinki. Furthermore, the study protocol and Informed Consent were approved in 2021 by the Ethics Committee of the Faculty of Medicine of the University of Coimbra (CE-060/2021). Based on previous research on neurofeedback (***Pereira et al., 2023***), we aimed for an effect size of 1.09 (*α* = 0.05) for a modulation activity difference between active and sham conditions. Based on G*Power (***Faul et al., 2007***), the estimated sample size per group was 19 participants.

Participants had an average mini-PROMS score of 19.3 ± 5.6 (range: 11.0-31.5), indicating their level of musical ability. Motor imagery ability scores were 5.8 ± 0.7 for internal visual imagery (range: 4.3-7.0), 5.6 ± 1.1 for external visual imagery (range: 3.0-7.0), and 5.4 ± 0.9 for kinesthetic imagery (range: 4.0-7.0). All participants were right-handed, with an average laterality index of 83.8 ± 13.6 (range: 45.0-100.0).

### Paradigms

The main objective of the functional localizer was to identify the brain networks of motor imagery and music interpretation. The trial design is presented in Figure 3A, comprised of 6 trials, each containing 6 seconds of ‘Rest’, 10 seconds of ‘Motor imagery’, ‘Music’, and ‘Noise’, and 6 seconds of ‘Reward report’ conditions, and with a total duration of approximately 7 minutes. This run was used to identify the premotor cortex on the left and right hemispheres during the NF session, which are the target regions for the NF imagery runs. The participants were asked to imagine tapping the fingers of their hands during the ‘Motor imagery’ condition and to attentively listen to the music during the ‘Music’ condition. Afterward, they were asked to report the pleasantness of the music in five levels (unpleasant, slightly unpleasant, neutral, slightly pleasant, pleasant). The music played was sampled from the chords that were used in the imagery runs as feedback. In this run, the instructions and rating scale appeared on the screen.

**Figure 3.**
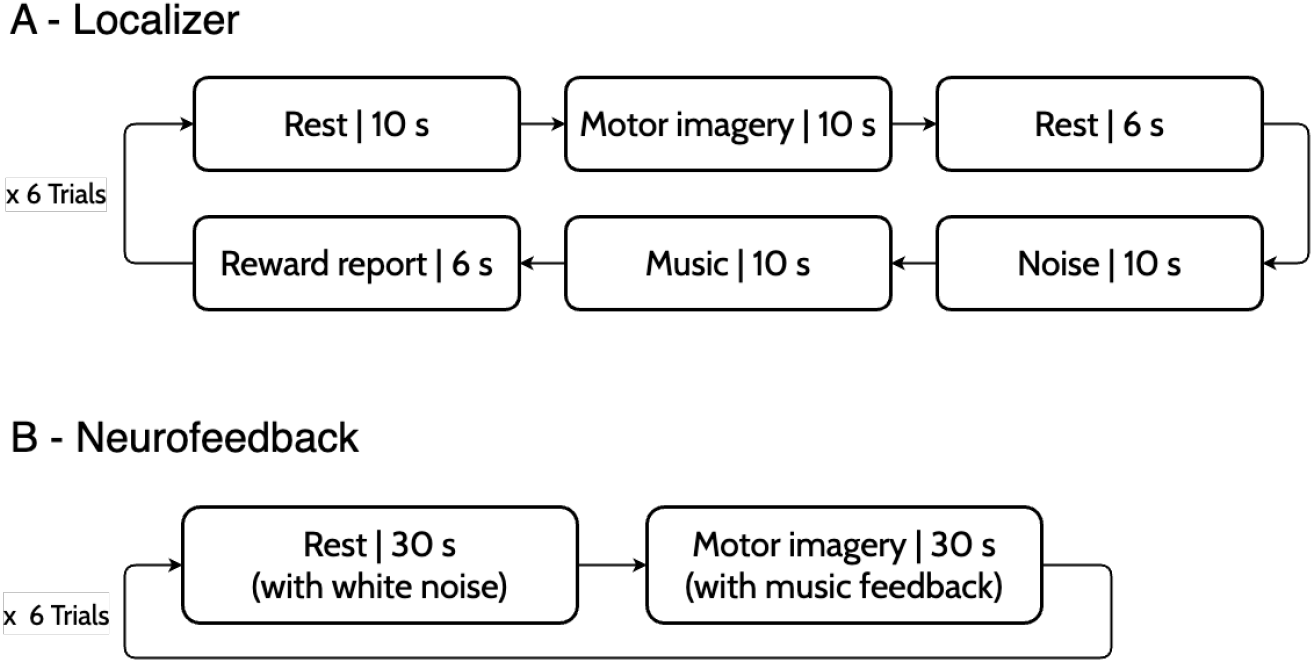
Functional paradigms for the localizer (A) and neurofeedback (B) runs. The localizer comprised 6 trials of interleaved blocks of rest, motor imagery, music listening, and music pleasantness report. During the NF runs, participants trained to control their PMC’s interhemispheric connectivity during the ‘Motor imagery’ blocks while imagining their hands and fingers moving. The feedback, presented only during these blocks, is based on a pre-validated rewarding chord progression that evolves or regresses according to the windowed correlation between the activities of the left and right PMCs.

The main objective of the NF runs was to train the participants in modulating the interhemispheric connectivity of the PMC through the imagery of motor actions. The trial design is presented in Figure 3B. The ‘Motor imagery’ was accompanied by musical feedback, which was calculated using the windowed correlation (8-second window) between the activity of the left and right premotor cortices. Participants were asked to imagine moving their hands and fingers (without actually moving) while listening to the music feedback. The instruction was auditory the ‘Rest’ condition was indicated by white noise, and the ‘Motor imagery’ condition by the music feedback. Following the task of (***Pereira et al., 2019***), halfway through the 30-second ‘Motor imagery’ block, a ‘beep’ sound was played - this indicated that participants should slowly decrease the frequency of the imagined movements until stopping close to the end of the block. This allowed for a prolonged variation of the BOLD activity in both premotor cortices, which would lead to a prolonged high correlation throughout the full block.

### Neurofeedback Setup

#### Feedback Music

In our approach, we mapped the correlation values to a harmonic progression where each 0.1 increment in correlation corresponded to a specific base note, allowing for ten possible base notes. The harmonic progression was dynamically shaped by these base notes, with the level of correlation (ranging from 0 to 1) determining the tonal foundation. Furthermore, the chord type was influenced by correlation changes over time: when the correlation increased compared to the previous time window, the chord was either major or major 7th, whereas when the correlation decreased, a diminished 7th chord was played. This real-time mapping ensured that musical pleasantness, and hence positive or negative feedback, fluctuated in alignment with the correlation variation. In Figure 4, we illustrate a chord progression derived from a correlation time course, showing how each time point was associated with the chord the participant heard. The resulting harmonic shifts reflect the programmed relationship between correlation and musical feedback. We provide the audio of this example as supplementary material.

**Figure 4.**
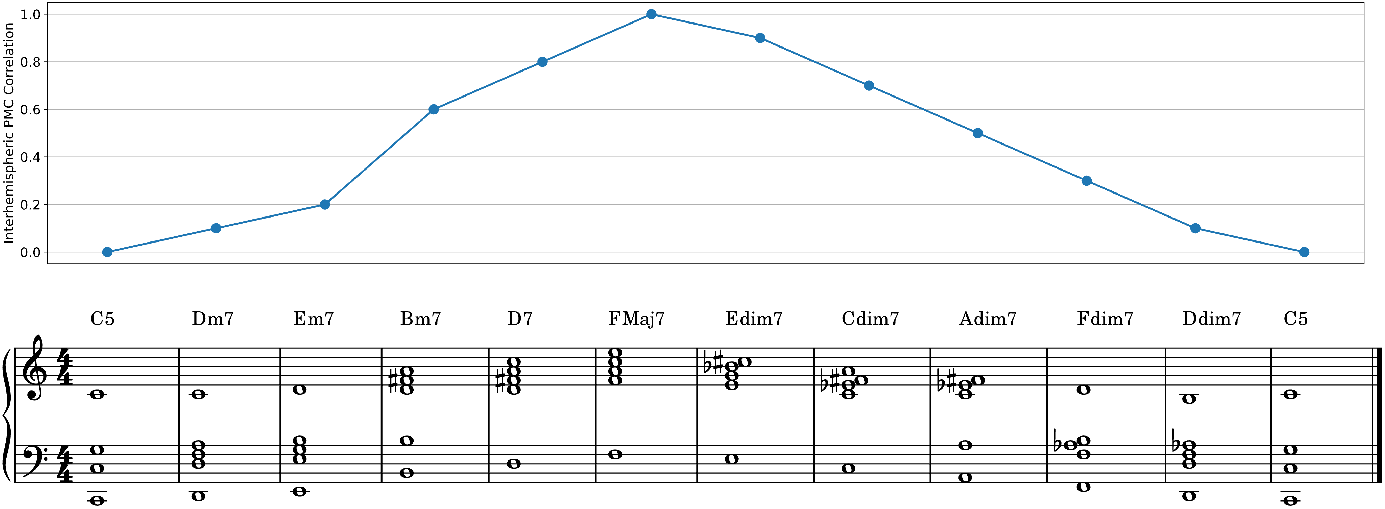
Feedback music - correspondence between an example interhemispheric brain correlation time course and its translation into musical chords/feedback presented. Positive feedback is linked to higher base notes and more pleasant chords (major, major 7th), while negative feedback is linked to lower base notes and less pleasant chords (diminished 7th).

#### Setup at the MRI

The technical neurofeedback loop used during the MRI session is illustrated in Figure 5. The MRI console exported DICOM files in real-time, i.e., as they were acquired, using a proprietary addin that transmitted the data to the real-time processing computer. This computer ran Turbo-BrainVoyager v4 software for real-time fMRI analysis, where the target regions of interest (ROIs) were previously defined in the bilateral primary motor cortex (PMC) of each participant based on anatomical landmarks and functional activation observed during the localizer run (‘Motor imagery’ vs. ‘Rest’). Inter-run alignment was activated, which ensured the ROIs stayed valid during the session. A separate feedback monitoring computer ran a custom Python script that either received the windowed correlation values via TCP/IP for the real NF runs or generated random feedback for the sham NF runs, converted these values into a scale, and generated the MIDI signals which were sent to virtual instruments created in Logic Pro v11.1.2. The resulting audio feedback was delivered to the participants, updated every TR, through the noise-canceling headphones.

**Figure 5.**
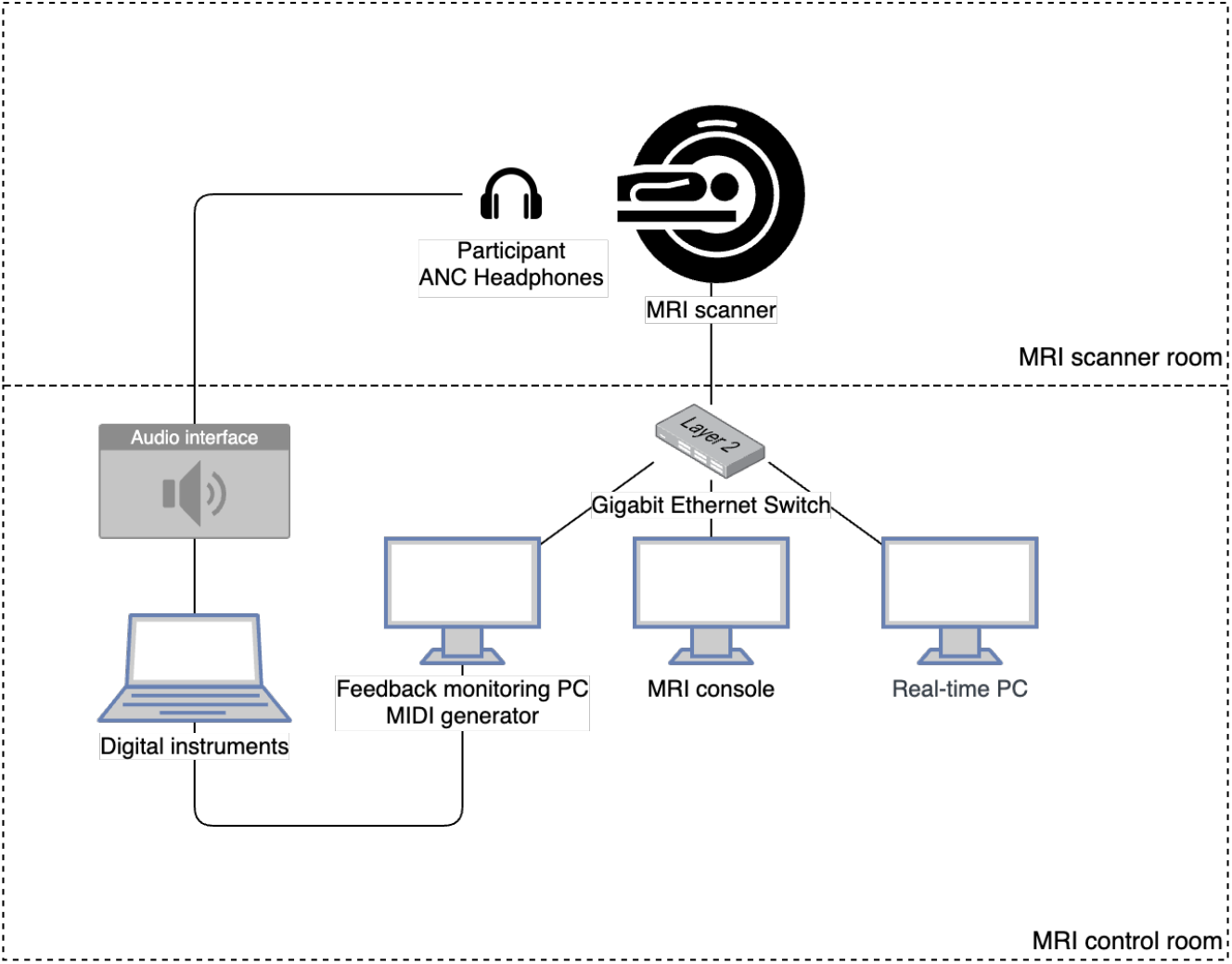
Schematic diagram of the real-time fMRI setup used to deliver music-based neurofeedback. The system enables real-time computation of interhemispheric correlation from fMRI signals and its translation into auditory feedback via MIDI generation. Functional images are acquired by the MRI scanner and processed by a real-time PC, with results transmitted over a local network to the feedback monitoring PC. This PC maps correlation values to musical parameters, generating MIDI signals that are converted to sound by digital instruments and played to the participant through active noise-cancelling (ANC) headphones.

#### Real-time analysis

Real-time analysis was performed in Turbo-BrainVoyager v4. After the localizer run, the ROIs in the bilateral premotor cortices of each participant were defined based on the activation map of the contrast between ‘Motor imagery’ and ‘Rest’ conditions. The statistical threshold was set to t-value > 3.1 (p < 0.001 uncorrected). Anatomical landmarks of the precentral gyrus, visible when superimposing the anatomical T1w image, were also considered in this definition. For all functional runs, the preprocessing steps included motion correction, inter-run alignment, and detrending. An iterative General Linear Model (GLM) was used to estimate the activation maps for each run. The design matrices included predictors for each of the experimental conditions and the six motion parameters (translation and rotation in the X, Y, and Z axes) as confound predictors.

### MRI acquisition

The MRI acquisition was performed with a 3 T Siemens Magnetom Prisma fit scanner with a 64-channel head coil at the Institute of Nuclear Sciences Applied to Health from the University of Coimbra. Auditory stimuli were presented using MRI-compatible headphones (Optoacoustics Optoactive II). The scanning session started with the acquisition of two anatomical images: one 3D anatomical magnetization-prepared rapid acquisition gradient echo pulse sequence (repetition time (TR) = 2530 ms, echo time (TE) = 3.5 ms, flip angle (FA) = 7°, 176 slices, voxel size 1.0 × 1.0 × 1.0 mm, field of view (FOV) = 256 × 256 mm) and one T2 space sequence (TR = 3200 ms, TE = 410 ms, 176 slices, voxel size 1.0 × 1.0 × 1.0 mm, FOV = 256 mm). A total of five functional runs were acquired using a 2D gradient-echo echo-planar imaging sequence (TR = 1500 ms, TE = 30 ms, flip angle = 75°, 26 slices, voxel size = 3.0 × 3.0 × 3.5 mm, slice gap = 0.5 mm, FOV = 210 mm, GRAPPA factor 2, echo spacing = 0.67 ms, bandwidth = 1700 Hz/px). For unwarping, a gradient field mapping sequence was acquired before the functional scans (TR = 400 ms, TE1 = 4.92 ms, TE2 = 7.38 ms, flip angle = 60°, 26 slices, voxel size = 3.0 × 3.0 × 3.5 mm, slice gap = 0.5 mm, FOV = 210 mm, bandwidth = 600 Hz/px). The participants’ physiological signals (respiratory and cardiac) were recorded during the functional runs using the scanner’s Physiological Measurement Unit (PMU). The respiratory signal was recorded at 50 Hz using a respiratory cushion, and the cardiac cycle was recorded at 200 Hz using a pulse sensor.

### Offline MRI analysis

The MRI data were organized according to the Brain Imaging Data Standard (BIDS), using a customizable conversion tool (***Boré et al., 2023***) for the DICOM images and custom scripts to associate the task events. Preprocessing was performed in fMRIPrep v24.0.1 (***Esteban et al., 2019, 2023***) and included slice time correction, motion correction, unwarping, and normalization to the MNI space. For a complete description of the fMRIPrep methods, please refer to the Supplementary Materials. All subsequent analyses were performed using custom Python scripts based on the package Nilearn v0.10.4 (***Abraham et al., 2014***;***Nilearn contributors et al., 2025***).

#### Visual interface dataset processing

To compare our music-based interface with the visual interface used in (***Pereira et al., 2019***), we obtained raw data previously acquired by the authors. The dataset was then converted to the BIDS format, preprocessed using the same version of fMRIPrep, and analyzed with an identical statistical approach based on GLMs, as the task is identical, only differing in the feedback modality. For a detailed description of the acquisition parameters and protocol, please refer to the original paper (***Pereira et al., 2019***).

### Statistical analysis

#### Behavioral data

During the localizer run, participants reported how pleasant/unpleasant each chord was to listen to: they rated each chord on a 5-level scale from unpleasant to pleasant, using the available buttons inside the MRI. We aimed to confirm that the pleasant and unpleasant labels we defined were valid for our sample. After extracting the report values for each type of chord, we statistically compared pleasant and unpleasant chords using a Mann-Whitney U test.

Participants answered the POMS questionnaire immediately before and after the MRI neuro-feedback session. The statistical comparison between the pre- and post-session POMS scores was performed using a Wilcoxon signed-rank test (paired samples) with Holm-Bonferroni correction (p=0.05).

We also look for correlations between the behavioral metrics (POMS, Mini-PROMS, MIQ-3) and the success metric using linear regression.

#### Activation maps

For the first-level analyses, GLMs were employed to obtain activation maps for each of the contrasts of interest and participant. For the localizer run, we estimated the contrast ‘Music’ + ‘Motor imagery’ > ‘Noise’ + ‘Rest’, while for the NF runs, the contrast was ‘Motor imagery’ > ‘Rest’. The design matrices included predictors for each of the experimental conditions and confound predictors for head motion (six motion parameters + first-order derivatives + powers, a total of 24), the mean signals of the cerebro-spinal fluid (CSF) and white matter masks, and 20 regressors based on the physiological signals, including RETROICOR, respiratory volume per time and heart rate variability responses (RVT/HRV), estimated with PhysIO toolbox (***Kasper et al., 2017***) - these last are critical when denoising data from connectivity-based NF experiments (***Weiss et al., 2020***). We also applied temporal high-pass filtering with a cut-off frequency of 0.008 Hz and considered a second-order autoregressive model AR(2) as the temporal variance model. The second-level group analyses considered spatial smoothing with a Gaussian kernel of full-width at half maximum = 6 mm and were corrected with False Discovery Rate (FDR) at q=0.005 (localizer, single run per participant) or with Bonferroni’s method at p=0.05 (NF runs, two per participant and feedback type). Additionally, we considered a minimum cluster size of 10 voxels.

#### Comparing active and sham feedback

One of the questions of this feasibility study regards the activation patterns of contingent feedback vs. sham. For this comparison, we considered the thresholded group-level activation maps (p=0.05 with Bonferroni’s correction, k>10) of active and sham feedback runs and defined a binary mask including the significant voxels for each of them. Then, we summed these masks to obtain a map of overlap between the two conditions - in practice, in each voxel, we show if the activity was significant for both conditions, only for active, or only for sham feedback. This map provides an overview of the pattern of network recruitment, while we look more closely at the reward and learning circuits.

#### Comparing music and visual interfaces

To search for differences in the activation maps retrieved from the visual interface experiment and our music-based active NF runs, we extracted the contrast maps between ‘Motor imagery’ and ‘Rest’ for all subjects (N=22 for music and N=10 for visual interfaces). Then, we employed a secondlevel, unpaired, two-sample t-test on the brain maps to see the effect of the interface difference across subjects. This comparison was corrected with Bonferroni’s method at p=0.05.

## Results

### Localizing the PMC for neurofeedback

The localizer run allowed for the functional definition of each participant’s left and right PMC, the targets for the neurofeedback runs, during the MRI session. This step was performed online based on the activation map contrasting the response to the ‘Motor imagery’ condition and the ‘Rest’ condition. The average MNI coordinates of the left PMC were (−36, −6, 54) and for the right PMC (37, −4, 55). The coordinates for each participant’s target regions are provided in Supplementary Table S1. The map showing the percentage of overlap of these regions across participants is presented in Figure 6.

**Figure 6.**
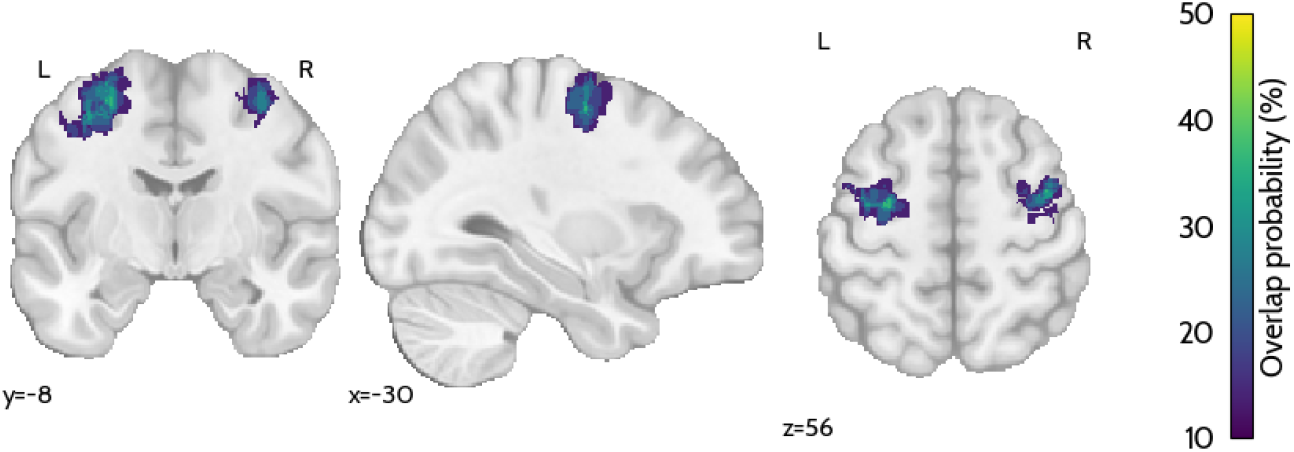
Brain probability map showing the percentage of overlap of the target ROIs (bilateral PMC) across participants.

Based on the localizer data, we look into the brain regions expected to be recruited during NF. In this test, we contrasted the activity both during motor imagery and music listening conditions against noise listening and rest conditions. The statistical map is shown in Figure 7. By performing clustering in this map, we identify significant clusters in the frontal inferior operculum, insula, precentral gyrus, supplementary motor area, temporal superior gyrus, Helschl’s gyrus, and cerebellum (clusters detailed in Supplementary Table S2). We also note the very clear deactivation of the default mode network, a marker of task engagement.

**Figure 7.**
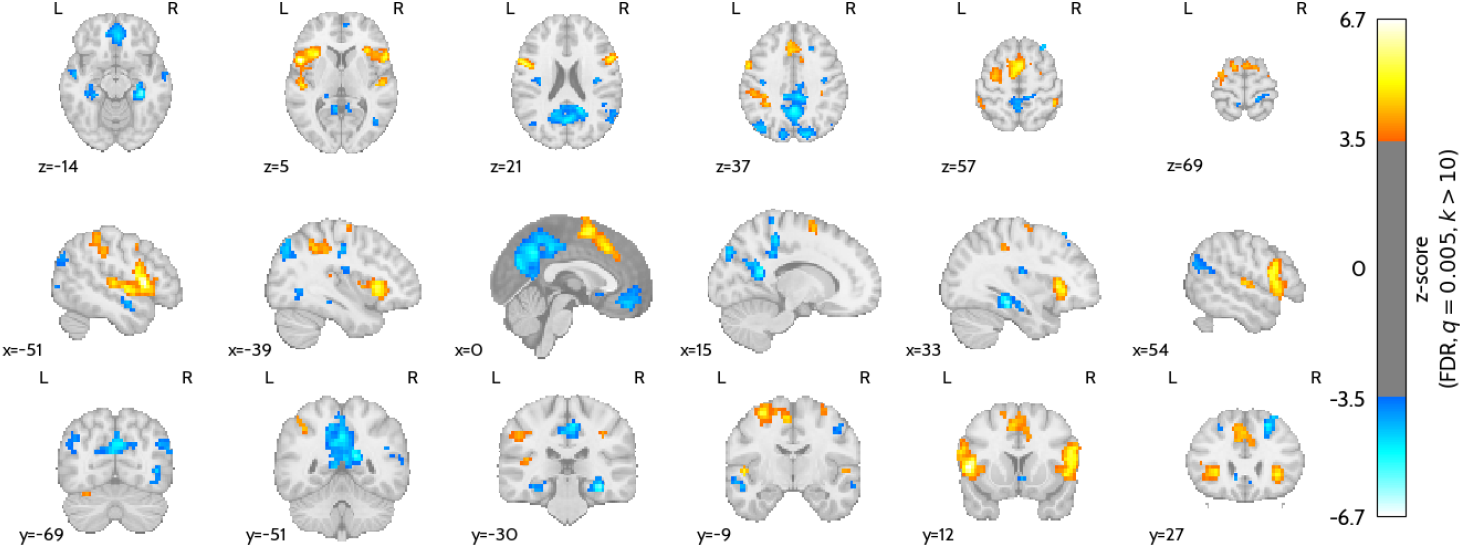
Brain activation map of the localizer run, contrasting motor imagery and music conditions vs. noise listening and rest. This map highlights regions linked to music perception (Helschl’s gyrus, temporal superior gyrus) and motor imagery (precentral gyrus, SMA).

The behavioral data from the reports regarding how pleasant/unpleasant each chord was to listen to confirmed our association (Supplementary Figure S3).

### Assessing the modulation of PMC interhemispheric connectivity

The basis for the feedback provided, and also the modulation target for the participants, was Pearson’s correlation between the BOLD signals of the left and right PMCs, as defined in the localizer run. These correlations were estimated in an 8-second sliding window across time during the acquisition. In Figure 8, we display the average of the estimated correlations across subjects for the active and sham runs during the NF ‘Motor imagery’ and ‘Rest’ blocks. In Figure 9, we display the average time course of such correlation in both NF conditions.

**Figure 8.**
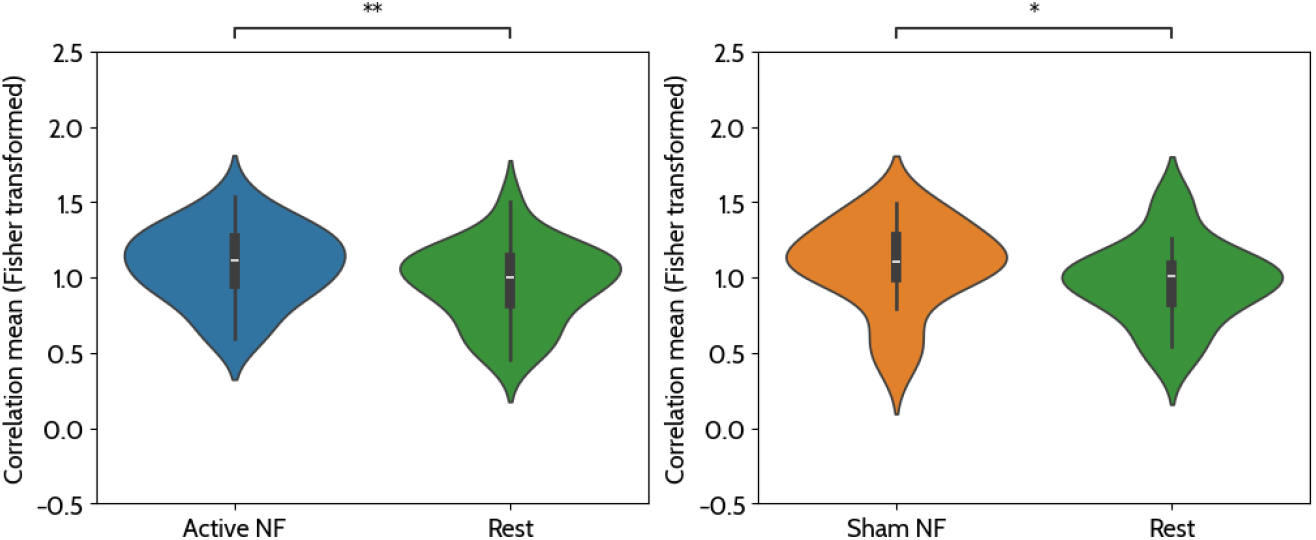
Average correlation values, as measured by the real-time software, during active NF, sham NF, and rest conditions across subjects (Fisher transformed). A one-sided Wilcoxon signed-rank test was conducted to compare mean correlation values between the NF and ‘rest’ conditions. Results showed that correlation values were significantly higher in the ‘active NF’ condition (Mdn = 1.11) compared to the ‘rest’ condition (Mdn = 1.00), W = 215.00, p = 0.003 (one-tailed, corrected for two comparisons) and also significantly higher in the ‘sham NF’ condition (Mdn = 1.11) compared to the ‘rest’ condition (Mdn = 1.00), W = 201.00, p = 0.014 (one-tailed, corrected for two comparisons).

**Figure 9.**
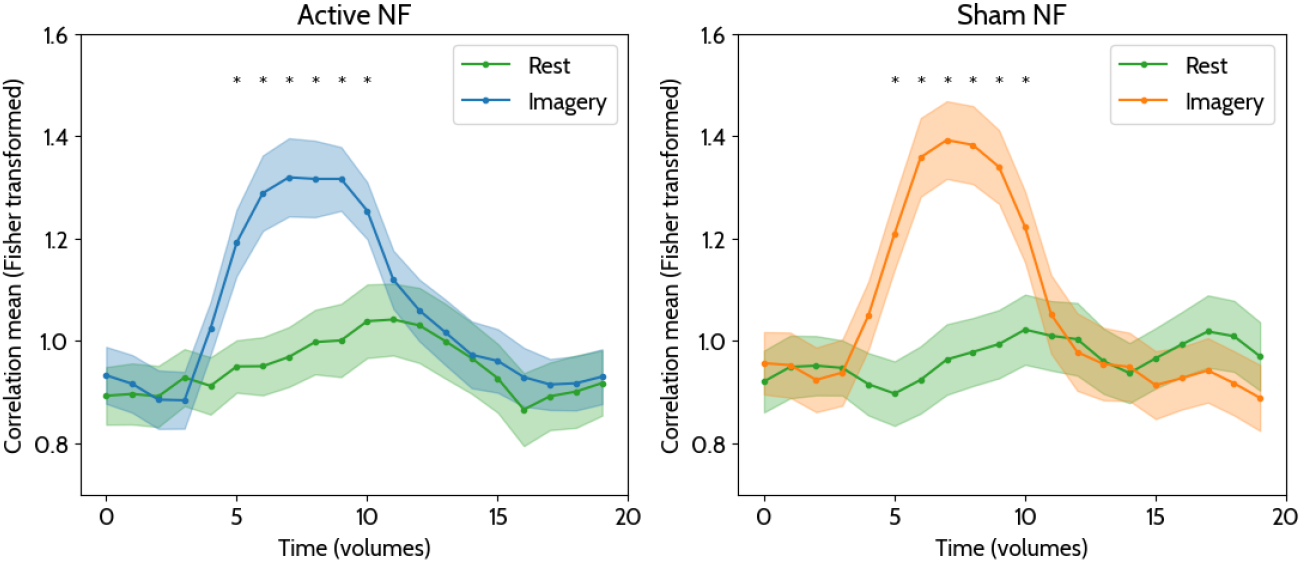
Average correlation time course during the ‘Motor imagery’ and ‘Rest’ conditions, as measured by the real-time software, for the active and sham NF runs. Each data point corresponds to the correlation in the sliding window that includes the 8 points that precede it. The correlation values were transformed to Fisher’s z. Paired Wilcoxon tests per timepoint between imagery and rest correlation values (* - p<0.05, corrected for two comparisons).

### Participant’s mood before and after the NF session

The assessment of participants’ mood before and after the MRI NF session was performed based on the self-reports of the POMS questionnaire. A significant decrease was found for the Tension sub-scale score (Mdn before = 3, Mdn after = 0, p = 0.012, W = 4) and for the Total score (Mdn before = 104, Mdn after = 98, p = 0.012, W = 28) - Figure 10. The results for the remaining sub-scales do not present a significant difference between time points and are presented in Figure S4.

**Figure 10.**
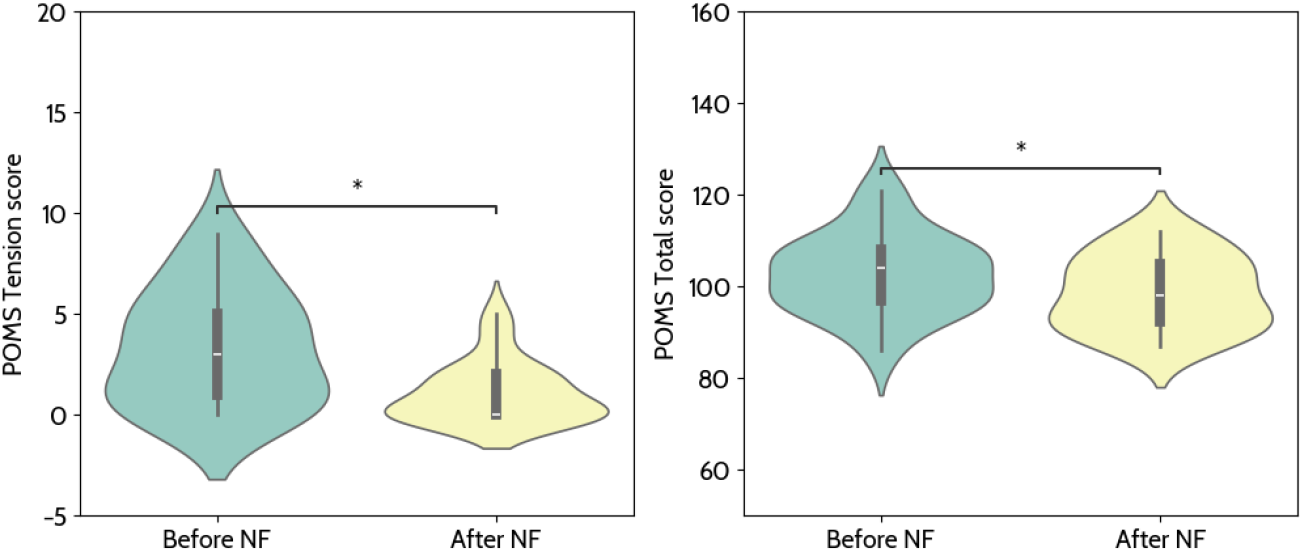
Comparison of the POMS scores before and after the NF session. We found a significant decrease in the tension subscale score and the total score with a paired sample Wilcoxon test after correcting for multiple comparisons with Bonferroni’s method (* - 0.01 < p < 0.05).

### Debriefing participants after the NF session

After the scanning session, participants were debriefed regarding the overall experience of the NF session, if they were able to perform the task, and lastly, if they could identify which was the best and worst run out of the four, both regarding their performance and the feedback contingency. 17 out of 22 participants (77%) reported an Active feedback run as the best, and 15 out of 22 participants (68%) reported a Sham feedback run as the worst. Also, most participants reported that imagery during the second part of the block (while decreasing imagined movement frequency) was harder when compared to the first part.

### Extracting brain activation patterns during neurofeedback

Figure 11 presents the brain activation maps for the active (A) and sham (B) neurofeedback runs, showing regions with significant activation when contrasting ‘Motor imagery’ with ‘Rest’. The corresponding clustering analysis (detailed in Supplementary Tables S3 and S4) reveals activation in several cortical and subcortical regions involved in motor planning, sensorimotor integration, auditory processing, and cognitive-affective regulation. Key regions include the supplementary motor area, precentral gyrus, and inferior parietal lobule, which are associated with motor execution and spatial coordination. Additionally, consistent engagement of the insula, putamen, and inferior frontal gyrus suggests involvement of interoceptive networks. The superior temporal gyrus and Heschl’s gyrus were also recruited, reflecting the role of auditory processing in the NF task.

**Figure 11.**
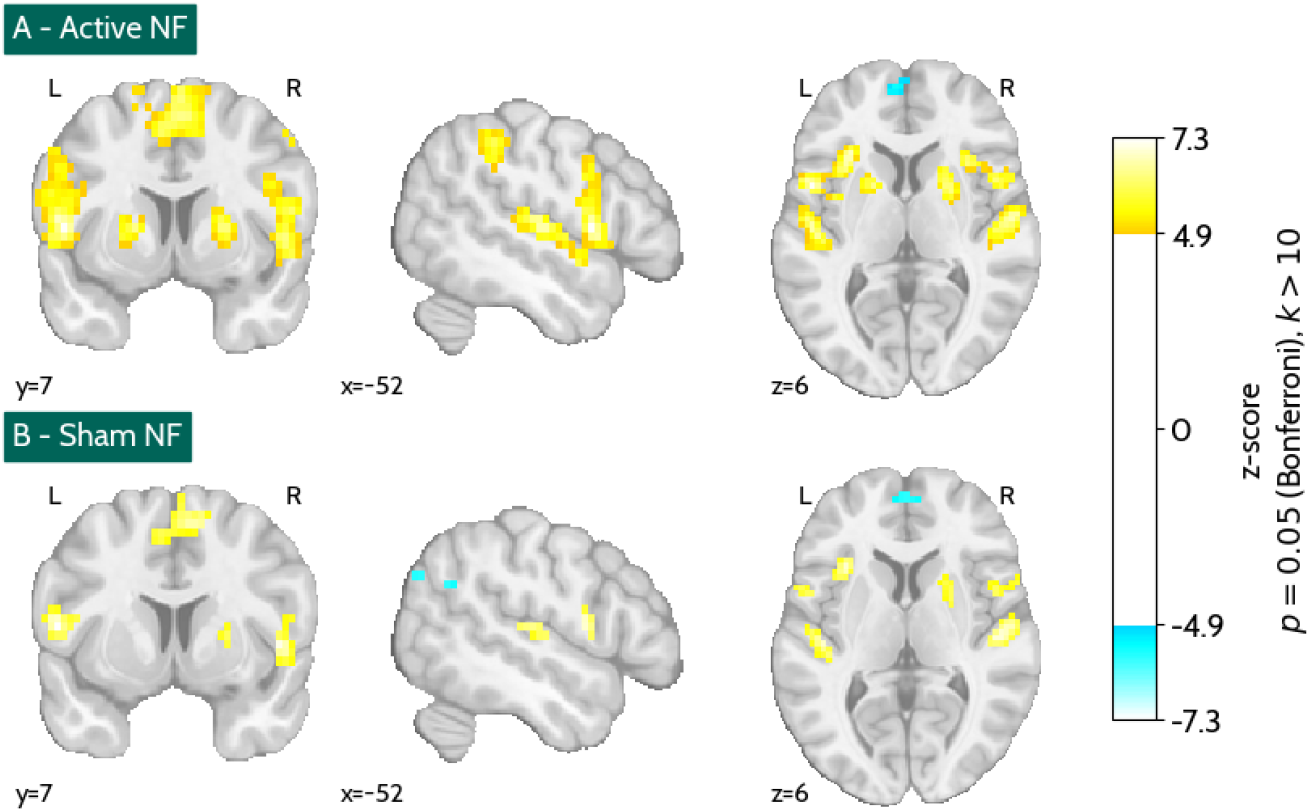
Brain activation maps of the active (A) and sham (B) neurofeedback runs contrasting ‘Motor imagery’ with ‘Rest’ (Bonferroni’s correction, p=0.05, k>10).

These regions were observed in both active and sham conditions, but with differential cluster peak activation and size. Figure 12 presents a comparison between the thresholded activation maps for the active and sham neurofeedback conditions, highlighting regions that exhibited significant activation exclusively in one condition or both. Table 1 provides a detailed breakdown of the clusters that were significantly active only in the active NF condition. These include regions such as the insula and putamen, precentral gyrus, supplementary motor area, inferior frontal gyrus, superior temporal gyrus, and inferior parietal lobule.

**Table 1.** Clustering of the active NF-only regions in Figure 12. For each cluster, we provide the coordinates in MNI space and the label of the AAL3 atlas.

| Cluster ID | X | Y | Z | AAL3 |
| --- | --- | --- | --- | --- |
| 1 | -57 | -27 | 10 | Temporal_Sup_L |
| 2 | -51 | -39 | 42 | Parietal_Inf_L |
| 3 | -48 | 6 | 6 | Frontal_Inf_Oper_L |
| 4 | -30 | -6 | 58 | Precentral_L |
| 5 | -30 | 24 | 2 | Insula_L |
| 6 | -21 | 6 | 6 | Putamen_L |
| 7 | -3 | 0 | 66 | Supp_Motor_Area_L |
| 8 | 9 | 6 | 66 | Supp_Motor_Area_R |
| 9 | 18 | 6 | 6 | Undefined |
| 10 | 27 | -6 | 50 | Precentral_R |
| 11 | 36 | 18 | 6 | Insula_R |
| 12 | 48 | 9 | 22 | Frontal_Inf_Oper_R |
| 13 | 48 | -24 | 14 | Temporal_Sup_R |
| 14 | 54 | 0 | 46 | Precentral_R |
| 15 | 54 | 3 | 2 | Rolandic_Oper_R |

**Figure 12.**
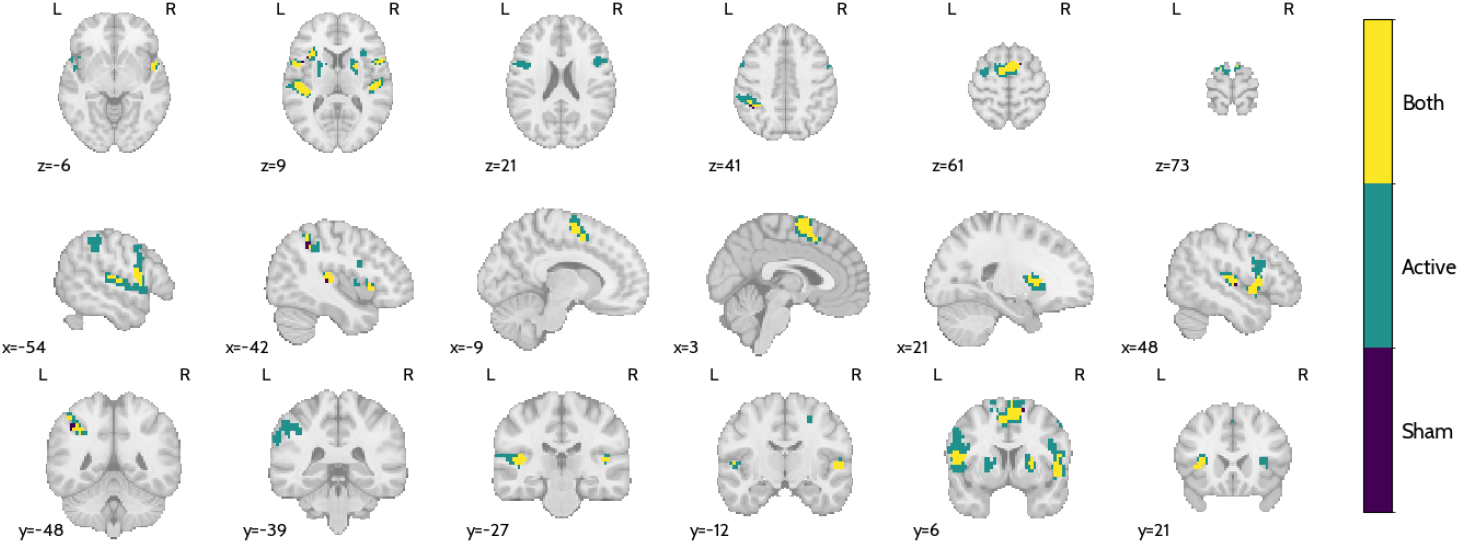
Comparison between the thresholded (Bonferroni’s correction, p=0.05, k>10) active and sham NF maps. Voxels for which both active and sham NF elicited significant activations are shown in yellow, while the ones for which only active NF elicited a significant activation are shown in green, and for sham-only in purple.

### Contrasting music and visual interfaces

Figure 13 presents the statistical contrast between the brain activity patterns elicited during music-based active neurofeedback and those observed in the visual-based active NF of (***Pereira et al., 2019***). Table S5 details the clusters where significant differences were found between these two datasets. Increased activation was observed in auditory-related regions, including Heschl’s gyrus, the superior temporal gyrus (bilaterally), and the Rolandic operculum, highlighting the engagement of auditory processing networks during music-based NF. Additionally, activation differences were identified in motor control-related areas, such as the SMA, precentral gyrus, insula, and putamen, suggesting potential differences in sensorimotor integration between the two feedback modalities. The mid-cingulate cortex and inferior frontal gyrus also showed differential activation, which may be linked to cognitive control and emotional processing.

**Figure 13.**
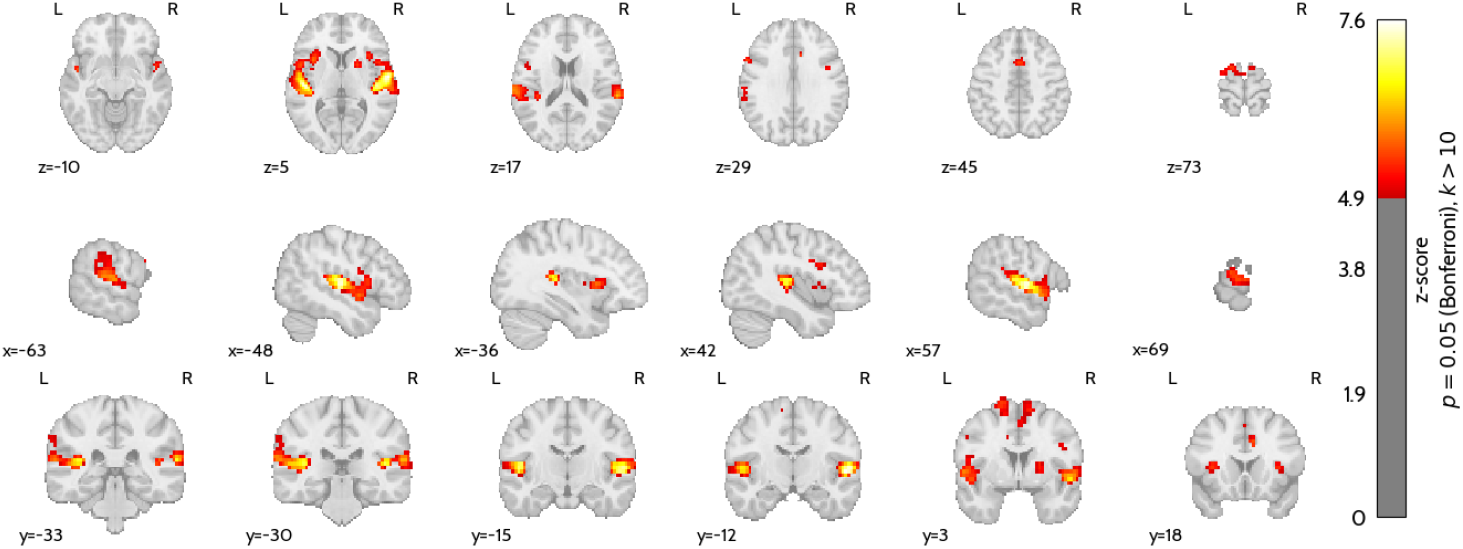
Brain map for the statistical contrast between the music-based active NF runs and the active NF runs with the visual interface of (***Pereira et al., 2019***) - ‘Motor imagery’ vs. ‘Rest’ (Bonferroni’s correction, p=0.05, k>10).

## Discussion

This study probed the use of music as a neurofeedback interface and investigated its capacity to modulate interhemispheric connectivity during motor imagery and the impact on affective measures such as mood. By comparing music-based feedback with both sham and visual neurofeedback conditions, we aimed to characterize the neural mechanisms engaged by different feedback modalities and assess the feasibility of using connectivity-based NF coupled with music as a neurobehavioral and impactful alternative to classical NF approaches.

### Modulation of PMC interhemispheric connectivity

The primary objective of neurofeedback training in this study was to modulate interhemispheric connectivity in the PMC using music feedback. Importantly, the determination of whether the training was effective is a matter of debate, with currently no standardized method for measuring success (***Paret et al., 2019***). Here, we defined the success metric as a significant difference in PMC correlation between ‘Motor imagery’ and ‘Rest’ conditions. Our results indicate that this modulation was successfully achieved, as the main correlation during imagery blocks was stronger than during rest.

This result was particularly evident for the first half of the block, a modulation pattern which was also observed in (***Pereira et al., 2019***). Sustaining correlation is not trivial, and actually requires skilled imagery, because as the BOLD signal reaches the plateau, correlation decreases to around zero this is why we instructed participants to change the frequency of the imagined movement after the middle of the block. However, we did not verify the hypothesized decrease in BOLD after the ‘beep’ to keep a high correlation (Supplementary Figure S1), which consistently matches participants’ reports about the difficulty of decreasing imagery frequency.

### The impact of music-based connectivity NF on behavioral measures and mood

Attentional factors seem to play a key role in both neurofeedback performance and learning, while motivation and mood have been identified as moderate predictors of success (***Kadosh and Staunton, 2019***). Here, we assessed changes in mood as a target outcome measure via the POMS self-reporting questionnaire. We found that the overall score and the Tension subscale decreased after the NF session, which are signs of mood improvement. This scale has been found to change after NF in previous studies reporting a full NF intervention in autism spectrum disorder using a visual interface exploiting facial expressions (***Direito et al., 2021***) and in major depressive disorder using a visual thermometer (***Mehler et al., 2018, 2021***). In our case, none of the differences are correlated with the success metric (Supplementary Figure S7).

Following up on these results, performing a music-based NF task could have a direct clinical effect, since it provides the opportunity for emotion control. We recently reviewed how music is being used in neurofeedback loops (***Sayal et al., 2025***), finding six studies out of the 15 reviewed also reporting significant changes in behavioral metrics (e.g., affective and mood-related measures, cognitive functioning scores, attention and mental control scores) after the NF intervention.

Regarding the Mini-PROMS and MIQ-3 questionnaires, we found no statistically significant correlations with the success metric (Figures S5 and S6). This means that, in this case, none of these would predict individual neurofeedback success (***Haugg et al., 2021***).

### Participants’ perception of contingency

The reported best and worst runs, as perceived by the participants, mostly match one of the active and sham runs, respectively. This reassures us that participants were able to perceive the presence of contingent, i.e., valid, feedback. We note that the participants were unaware that sham feedback was going to be used in two of the NF runs. The utilization of sham feedback as the control condition for NF specificity is a matter of debate (***Paret et al., 2019***). The detection of non-contingent feedback, i.e., the loss of control, could lead to frustration or the abandoning of a valid strategy of neuromodulation. Anecdotally, some participants reported that after a sham run, in which the feedback did not work as expected, they tried harder to focus on the imagery task and make the feedback work - this is a possible confounding effect that might partially explain the results that follow.

### Active vs. Sham feedback

In a sham-controlled study, the perception of whether the feedback signal aligns with effort may not differ significantly between groups (***Sorger et al., 2019***). Many participants receiving real neurofeedback struggle to perceive a direct relationship between their effort and signal changes due to the inherent difficulty of self-regulating neural activity (***Ros et al., 2013***; ***Thibault et al., 2016***). This disconnect between intention and perceived outcome contributes to a common experience of non-contingency, reported across both active and sham conditions (***Lubianiker et al., 2019***; ***Ninaus et al., 2013***). If participants apply consistent effort, the relationship between effort and signal change is likely to be of similar magnitude in both groups. This perceived equivalence in feedback success may balance participants’ subjective experience, despite differences in signal origin.

Our results support this interpretation: we found no significant difference in the neural success metric when comparing active and sham feedback conditions. This outcome was anticipated, but we hypothesized that differences would emerge in the broader network dynamics, particularly in reward and learning-related circuits, namely the putamen (***Dias et al., 2024***). This hypothesis is supported by previous work suggesting that contingent feedback engages reinforcement learning processes, even in the absence of conscious performance evaluation (***Sitaram et al., 2017, 2024***). Remarkably, we observed greater and more extensive activation in the putamen and insula during active NF. These regions have well-established roles in the context of NF: the putamen, as part of the basal ganglia, is involved in motor learning, reward prediction, and habit formation, while the insula is crucial for interoception, emotional salience, and monitoring internal states. Both are frequently reported in NF studies examining learning, motivation, and affective processing (***Emmert et al., 2016***; ***Sitaram et al., 2024***). Their stronger engagement in the active condition suggests that, although the success metric alone may not distinguish between groups, the contingency of the feedback loop in the active NF condition recruits additional neural systems, potentially supporting the development of internal models of control and reward-based learning.

### Music vs. visual interface for NF

Previous research has integrated music into NF paradigms, as reviewed in (***Sayal et al., 2025***), yet important questions remain. One key aspect of these implementations is the understanding of the neural correlates of music-evoked emotions, which have been extensively characterized (***Koelsch, 2020***). Music engages brain regions involved in emotion processing and reward, such as the limbic system, auditory cortex, and basal ganglia, overlapping with the networks typically targeted by neurofeedback (***Sitaram et al., 2024***). This overlap suggests that music may not only serve as a feedback modality but may also potentiate NF effects by co-activating these functional systems. In our results comparing brain activity during music-based and visual neurofeedback, we observed stronger activation in music-related regions, including in the Heschl’s gyrus and supplementary motor area (SMA), but also in the putamen, insula, and the cingulate cortex (the latter also part of the salience network). These regions play distinct yet interconnected roles relevant to neurofeedback. Heschl’s gyrus, part of the primary auditory cortex, is crucial for processing complex sound features, and its activation indicates that participants were effectively perceiving and engaging with the musical feedback. The SMA is involved in motor planning and imagery, which aligns with the task demands and suggests successful task engagement. The putamen, a component of the basal ganglia, is implicated in motor learning and reward processing, and its activation may reflect reinforcement mechanisms facilitated by the emotionally salient and rewarding nature of music. Lastly, the cingulate cortex, particularly its mid and anterior regions, is associated with attention, cognitive control, and emotional regulation - all central to self-regulation and learning during NF. Its recruitment, together with the insula, suggests an important role for the saliency network.

Compared to traditional visual NF interfaces, which provide explicit and structured feedback, music offers a dynamic and emotionally engaging alternative. Music can act as a probe for emotional and reward-related processes, potentially enhancing NF learning through mechanisms such as emotional contagion (***Schubert, 2013***). This interplay between emotion, reward, and self-regulation may offer unique advantages over purely visual feedback, making music-based NF a promising avenue for further research.

### Future directions

In our study, we employed a harmonically simple but easy-to-understand chord progression as the interface to provide musical feedback of a motor imagery task. We manipulated the base note, translating the strength of connectivity, and what we defined as the ‘pleasantness’ of the chords (for the direction of change in correlation). This pleasantness can be understood as a form of musical valence, a key dimension for triggering positive vs. negative feedback, which has well-documented neural correlates when encoded in music (***Concina et al., 2019***; ***Goodkind et al., 2012***; ***Lindquist et al., 2016***; ***Sayal et al., 2024***).

Looking ahead, future studies could adopt a more naturalistic approach by integrating real music stimuli or more complex harmonic progressions. While this could enhance the ecological validity of music-based neurofeedback, it also introduces greater variability in how participants interpret the feedback. Balancing complexity and interpretability will be crucial in refining this approach for both experimental and clinical applications.

Beyond methodological advancements and a better understanding of the underlying neuroscience of music, we believe that the novel NF implementation presented here holds significant potential for clinical applications, particularly in emotion regulation, given the recruitment of regions of the salience network (***Pinto et al., 2023***), and in pain management, given the considerable overlap of the pain and music perception networks (***Sihvonen et al., 2022***). The effects of music on pain perception and its reduction are well-documented (***Arnold et al., 2024***), yet their clinical relevance remains underexplored (***Sihvonen et al., 2022***; ***Werner et al., 2023***). Building on our previous work demonstrating the feasibility of NF in the context of pain empathy using a visual interface (***Travassos et al., 2020***), future clinical trials integrating this music-based NF interface could offer a novel framework to investigate the mechanisms of music-induced analgesia and its neural correlates, potentially paving the way for innovative therapeutic interventions.

## Supporting information

CRED-NF Checklist

Sample Chord Sequence Audio

## Acknowledgments

A word of gratitude to the MRI technicians at ICNAS, Sónia Afonso, Tânia Lopes, and Maria Inês Ferreira, for their help in designing the acquisition protocol and for their support during the neurofeedback acquisitions. We would like to thank all our participants for the time and dedication invested while volunteering for this study.

## Data and code availability

All code used in this study is available in the GitHub repository (https://github.com/CIBIT-UC/musicnf-novelinterface). The dataset, formatted in BIDS, can be accessed at Zenodo (https://doi.org/10.5281/zenodo.14803374). This study was preregistered on OSF (https://doi.org/10.17605/OSF.IO/AHXNB).

## Contributions

Conceptualization: AS, BD, TS, MCB; Data curation: AS, JP; Formal analysis: AS, BD, TS; Funding acquisition: BD, MCB; Investigation: AS; Methodology: AS, BD, TS; Resources: BD, MCB; Software: AS; Supervision: BD, TS, MCB; Validation: AS, JP, BD, TS, MCB; Visualization: AS, BD, TS; Writing - original draft: AS, BD, TS; Writing - review & editing: AS, JP, BD, TS, MCB.

