## Supplementary Materials for "A novel music-based real-time fMRI neurofeedback interface modulates interhemispheric connectivity and enhances mood"

#### 1. Localizer results

*Table S1 - Target ROIs coordinates in MNI of the left and right premotor cortices of each participant.*

| | Left | | | | Right | | | |
|---|---|---|---|---|---|---|---|---|
| ID | X | Y | Z | N voxels | X | Y | X | N voxels |
| 1 | -32.71 | -7.8 | 49.86 | 737 | 40.45 | -2.09 | 53.71 | 831 |
| 2 | -41.09 | -1.95 | 43.54 | 708 | 37.17 | -1.1 | 45.19 | 1348 |
| 3 | -44.21 | -1.59 | 45.97 | 1727 | 50.21 | -3.82 | 45.83 | 420 |
| 4 | -40.2 | -6.01 | 46.4 | 887 | 48.4 | -0.28 | 46.93 | 1572 |
| 5 | -28.42 | -2.08 | 63.03 | 3214 | 31.13 | 1.99 | 64.33 | 2088 |
| 6 | -32.56 | -8.62 | 42.41 | 2220 | 21.53 | -6.6 | 40.04 | 1613 |
| 7 | -38.91 | -7 | 42.81 | 1110 | 19.92 | -3.23 | 47.69 | 1235 |
| 8 | -49.45 | 0.6 | 51.12 | 2673 | 43.87 | -0.9 | 54.76 | 1762 |
| 9 | -47.68 | 1.43 | 48.15 | 1271 | 57.87 | 10.21 | 34.36 | 1475 |
| 10 | -31.52 | -4.88 | 55.3 | 2619 | 34.69 | -3.37 | 56.44 | 3013 |
| 11 | -21.97 | -9.77 | 60.11 | 1960 | 24.89 | -8.78 | 65.23 | 1861 |
| 12 | -40.68 | -1.17 | 70.49 | 1591 | 45.69 | -0.77 | 73.53 | 1459 |
| 13 | -37.86 | -7.12 | 61.21 | 1772 | 21.48 | -4.7 | 58.02 | 732 |
| 14 | -25.87 | -6.98 | 61.9 | 740 | 36.87 | -6.25 | 57.95 | 774 |
| 15 | -33.68 | -10.81 | 57.55 | 2285 | 37.15 | -11.82 | 60.17 | 1601 |
| 16 | -32.64 | -9.55 | 53 | 2718 | 43.32 | -6.5 | 56.84 | 1357 |
| 17 | -31.85 | -14.44 | 61.03 | 2170 | 35.57 | -16.49 | 61.57 | 2084 |
| 18 | -47.59 | -4.46 | 46.98 | 2121 | 42.35 | -1.8 | 53.72 | 1359 |
| 19 | -43.58 | -4.41 | 51.63 | 2153 | 39.43 | -6.02 | 51.12 | 1153 |
| 20 | -31.2 | -4.49 | 59.05 | 2657 | 31.92 | 0.25 | 60.59 | 1570 |
| 21 | -24.74 | -10.25 | 62.84 | 3112 | 35.84 | -11.33 | 60.32 | 2036 |
| 22 | -25.35 | -11.8 | 55.56 | 2386 | 31.08 | -6.92 | 56.12 | 1948 |
| Avg | -35.63 | -6.05 | 54.09 | 1947 | 36.86 | -4.11 | 54.75 | 1513 |
| Std | 7.83 | 4.09 | 7.63 | 753 | 9.47 | 5.36 | 8.67 | 551 |

*Table S2 - Clusters identified from the localizer activation map of Figure 7. MNI coordinates, peak z-value, cluster size in mm³, and labels from the AAL3 atlas.*

| Cluster ID | X | Y | Z | Peak z-value | Cluster Size (mm3) | AAL3 label |
|---|---|---|---|---|---|---|
| 1 | -51 | 12 | 6 | 6.74 | 20325 | Frontal_Inf_Oper_L |
| 1a | -54 | 6 | 22 | 5.81 | | Precentral_L |
| 1b | -39 | 18 | 2 | 5.57 | | Insula_L |
| 1c | -51 | -6 | -2 | 5.30 | | Temporal_Sup_L |
| 2 | 33 | 21 | 2 | 5.50 | 12568 | Undefined |
| 2a | 51 | 6 | 18 | 5.46 | | Frontal_Inf_Oper_R |
| 2b | 57 | 15 | 6 | 5.29 | | Frontal_Inf_Oper_R |
| 2c | 48 | 9 | -2 | 5.23 | | Insula_R |
| 3 | 3 | 3 | 62 | 5.40 | 22084 | Supp_Motor_Area_R |
| 3a | -33 | -9 | 62 | 5.32 | | Precentral_L |
| 3b | -3 | 21 | 42 | 5.17 | | Frontal_Sup_Medial_L |
| 3c | -6 | 6 | 54 | 5.12 | | Supp_Motor_Area_L |
| 4 | -36 | -48 | 42 | 5.40 | 6212 | Parietal_Inf_L |
| 4a | -48 | -36 | 50 | 4.50 | | Parietal_Inf_L |
| 4b | -36 | -33 | 38 | 4.49 | | Parietal_Inf_L |
| 4c | -51 | -33 | 38 | 4.30 | | Parietal_Inf_L |
| 5 | 54 | -18 | 6 | 4.64 | 1256 | Temporal_Sup_R |
| 5a | 45 | -21 | 6 | 4.45 | | Heschl_R |
| 6 | 45 | -42 | 58 | 4.64 | 2154 | Parietal_Sup_R |
| 6a | 42 | -36 | 50 | 4.51 | | Parietal_Inf_R |
| 6b | 36 | -33 | 42 | 4.33 | | Postcentral_R |
| 6c | 42 | -42 | 42 | 3.98 | | SupraMarginal_R |
| 7 | -42 | 42 | -2 | 4.43 | 395 | Frontal_Inf_Tri_L |
| 8 | -57 | -39 | 34 | 4.20 | 646 | SupraMarginal_L |
| 8a | -63 | -39 | 26 | 3.82 | | SupraMarginal_L |
| 9 | -30 | -66 | -22 | 4.16 | 502 | Cerebellum_6_L |
| 10 | 30 | -12 | 66 | 4.01 | 610 | Frontal_Sup_2_R |

### 2. Modulation of the target region

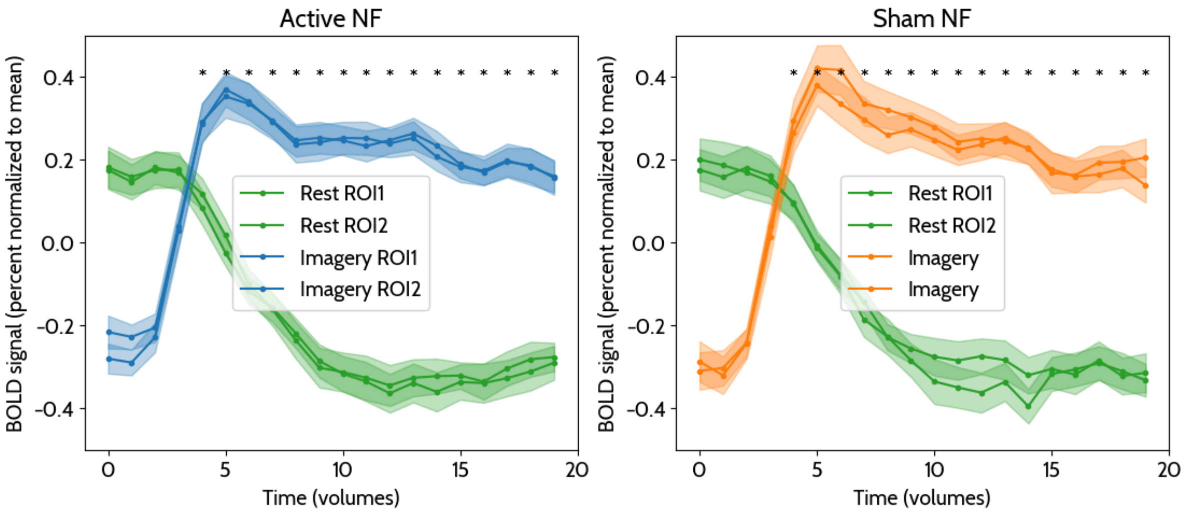


*Figure S1 - Average BOLD signals from the two target ROIs in the premotor cortex during imagery and rest conditions. A one-sided Wilcoxon signed-rank test was conducted to compare mean correlation values per timepoint between the 'Motor imagery' and 'Rest' conditions (one-tailed).*

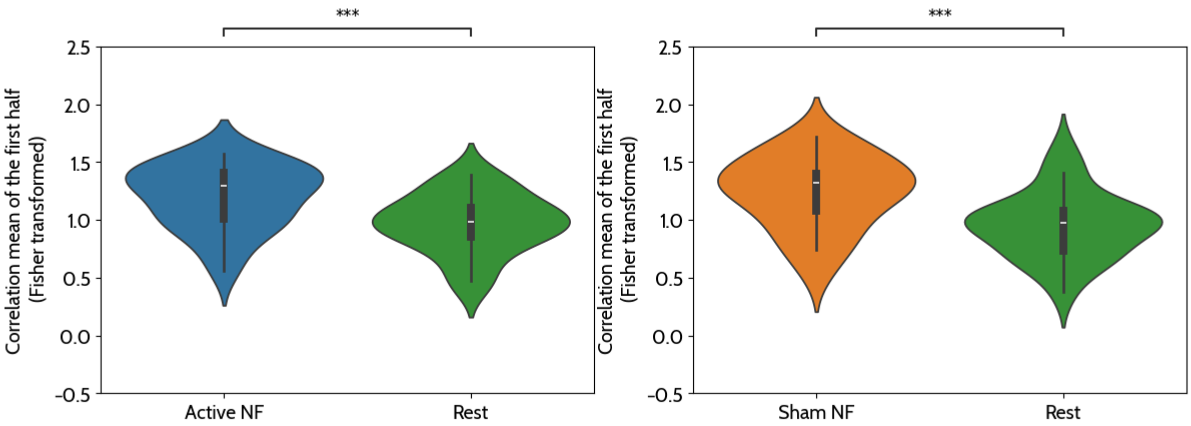


*Figure S2 - One-sided Wilcoxon signed-rank test was conducted to compare mean correlation values in the first half of the block between the 'active' and 'rest' conditions. Results showed that correlation values were significantly higher in the 'active' condition (Mdn = 1.30) compared to the 'rest' condition (Mdn = 0.98), W = 231.00, p = 0.000 (one-tailed) and significantly higher in the 'sham' condition (Mdn = 1.32) compared to the 'rest' condition (Mdn = 0.97), W = 231.00, p = 0.000 (one-tailed).*

### 3. Chord pleasantness validation

Reports were performed during the localizer run regarding how pleasant/unpleasant each chord was to listen to. Participants rated each chord on a 5-level scale from unpleasant to pleasant. The results are shown in Figure S3. The statistical comparison between pleasant and unpleasant chords was performed using a Mann-Whitney U test (two-sided, P_val:1.461e-17 U_stat=6.656e+03).

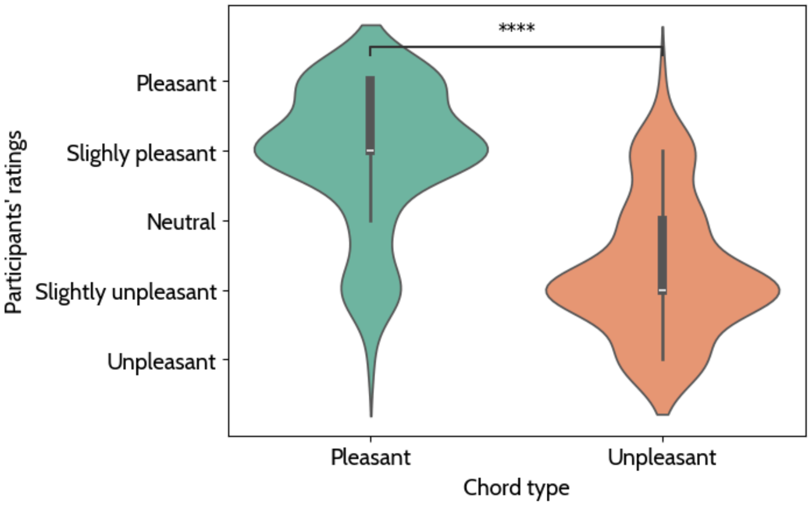


*Figure S3 - Participants' ratings of the pleasantness of each chord during the localizer run as violin plots.*

### 4. POMS subscales

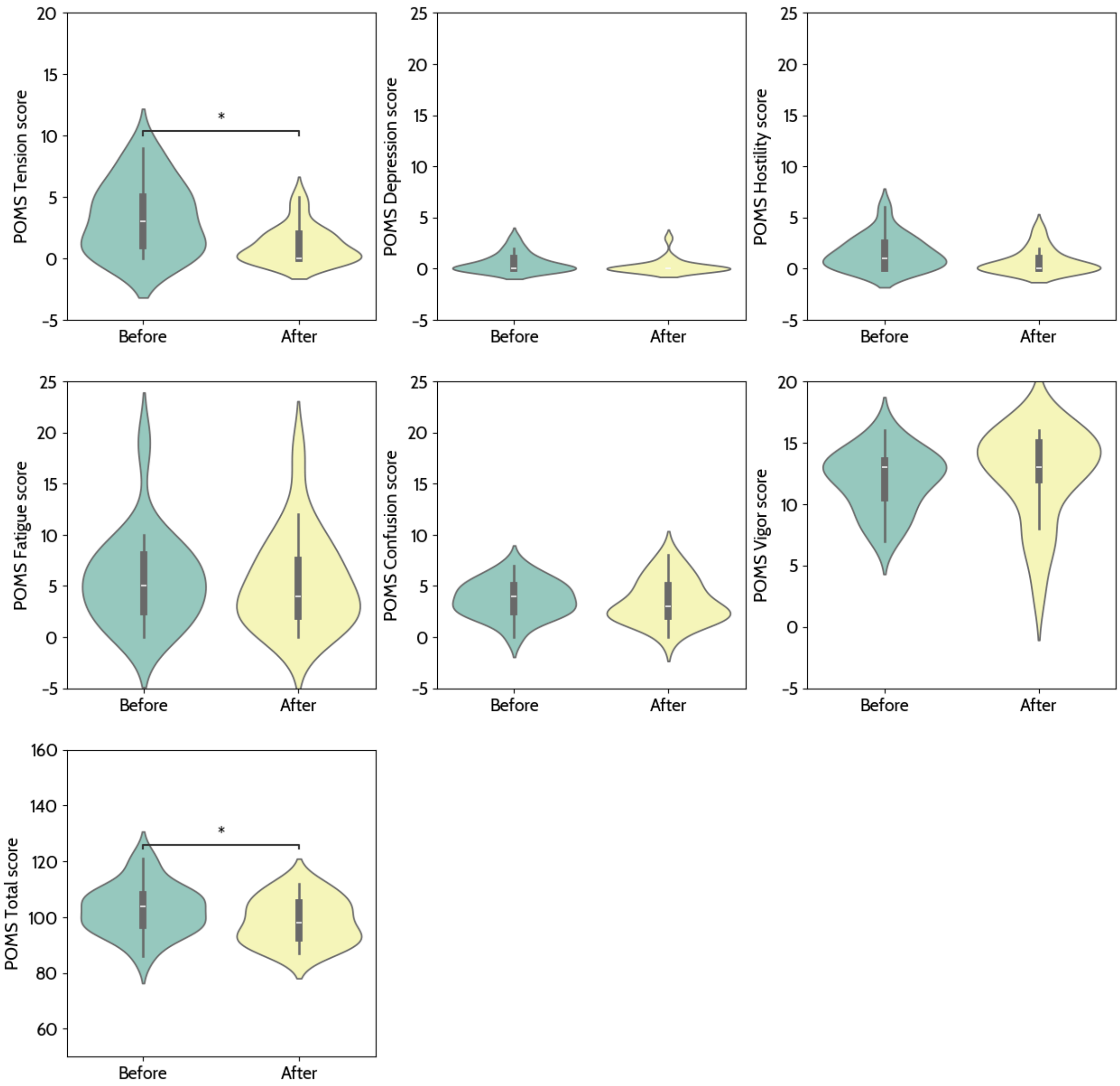


*Figure S4 - Statistical comparison between the POMS subscales before and after the intervention.*

### 5. Active and Sham activation clusters

*Table S3 - Table of clusters obtained from the activation map of Figure 11A, contrasting motor imagery with rest during active neurofeedback.*

| Cluster ID | X | Y | Z | Peak Stat | Cluster Size ($mm^3$) | AAL3 |
|---|---|---|---|---|---|---|
| 1 | -30 | 21 | 10 | 7.31 | 14256 | Insula_L |
| 1a | -51 | 6 | 6 | 7.25 | | Frontal_Inf_Oper_L |
| 1b | -51 | -15 | 6 | 6.68 | | Temporal_Sup_L |
| 1c | -48 | 0 | -10 | 6.06 | | Temporal_Sup_L |
| 2 | 57 | -12 | 6 | 6.70 | 10665 | Temporal_Sup_R |
| 2a | 51 | 6 | 2 | 6.35 | | Rolandic_Oper_R |
| 2b | 33 | 18 | 6 | 6.33 | | Insula_R |
| 2c | 51 | -3 | -2 | 6.31 | | Temporal_Sup_R |
| 3 | 0 | 3 | 58 | 6.63 | 10880 | Supp_Motor_Area_L |
| 3a | 3 | -3 | 66 | 6.59 | | Supp_Motor_Area_R |
| 3b | -18 | 3 | 70 | 6.06 | | Frontal_Sup_2_L |
| 4 | 27 | -9 | 50 | 6.57 | 1077 | Undefined |
| 5 | 21 | 15 | 2 | 6.50 | 2441 | Putamen_R |
| 5a | 24 | 3 | 6 | 6.34 | | Putamen_R |
| 6 | -18 | 6 | 6 | 6.43 | 1651 | Putamen_L |
| 6a | -24 | -3 | 14 | 5.81 | | Putamen_L |
| 7 | -54 | -39 | 46 | 6.05 | 5278 | Parietal_Inf_L |
| 7a | -63 | -36 | 38 | 5.67 | | Undefined |
| 7b | -63 | -39 | 30 | 5.56 | | SupraMarginal_L |
| 7c | -48 | -48 | 58 | 5.53 | | Parietal_Inf_L |
| 8 | -30 | -6 | 62 | 5.81 | 1580 | Precentral_L |
| 9 | 54 | 3 | 46 | 5.76 | 754 | Precentral_R |

*Table S4 - Table of clusters obtained from the activation map of Figure 11B, contrasting motor imagery with rest during sham neurofeedback.*

| Cluster ID | X | Y | Z | Peak Stat | Cluster Size ($mm^3$) | AAL3 |
|---|---|---|---|---|---|---|
| 1 | -54 | 6 | 10 | 6.59 | 1328 | Frontal_Inf_Oper_L |
| 2 | -42 | -27 | 10 | 6.52 | 1651 | Heschl_L |
| 3 | 48 | 3 | -2 | 6.40 | 4596 | Insula_R |
| 3a | 48 | -18 | 6 | 6.31 | | Heschl_R |
| 3b | 54 | -3 | -2 | 6.23 | | Temporal_Sup_R |
| 3c | 60 | 12 | 10 | 5.63 | | Frontal_Inf_Oper_R |
| 4 | 6 | 9 | 58 | 6.30 | 5278 | Supp_Motor_Area_R |
| 4a | -9 | -3 | 62 | 6.20 | | Supp_Motor_Area_L |
| 4b | 3 | 3 | 62 | 6.02 | | Supp_Motor_Area_R |
| 4c | -6 | 6 | 54 | 5.44 | | Supp_Motor_Area_L |
| 5 | -33 | 21 | 6 | 6.25 | 1220 | Insula_L |
| 6 | 24 | 3 | 10 | 5.58 | 718 | Putamen_R |
| 7 | -45 | -51 | 46 | 5.54 | 825 | Parietal_Inf_L |
| 7a | -36 | -48 | 42 | 5.41 | | Parietal_Inf_L |
| 7b | -48 | -42 | 42 | 5.18 | | Parietal_Inf_L |
| 7c | -45 | -51 | 54 | 5.16 | | Parietal_Inf_L |

### 6. Music vs. visual activation clusters

*Table S5 - Table of clusters obtained from the activation map of Figure 13, contrasting the visual and music NF interfaces.*

| Cluster ID | X | Y | Z | Peak Stat | Cluster Size ($mm^3$) | AAL3 |
|---|---|---|---|---|---|---|
| 1 | 60 | -9 | 6 | 7.58 | 17775 | Heschl_R |
| 1a | 51 | -24 | 10 | 7.28 | | Temporal_Sup_R |
| 1b | 54 | -3 | -2 | 7.28 | | Temporal_Sup_R |
| 1c | 63 | -33 | 14 | 6.73 | | Temporal_Sup_R |
| 2 | -51 | -18 | 10 | 7.44 | 22156 | Temporal_Sup_L |
| 2a | -39 | -30 | 10 | 6.82 | | Temporal_Sup_L |
| 2b | -51 | -6 | 2 | 6.56 | | Temporal_Sup_L |
| 2c | -57 | 3 | 2 | 5.90 | | Rolandic_Oper_L |
| 3 | 9 | 18 | 34 | 5.60 | 610 | Cingulate_Mid_R |
| 4 | -30 | -6 | 70 | 5.56 | 6966 | Frontal_Sup_2_L |
| 4a | 0 | 9 | 46 | 5.52 | | Supp_Motor_Area_L |
| 4b | -18 | 3 | 74 | 5.49 | | Undefined |
| 4c | 12 | 3 | 66 | 5.41 | | Supp_Motor_Area_R |
| 5 | -60 | 12 | 30 | 5.53 | 1041 | Precentral_L |
| 5a | -57 | 6 | 42 | 5.19 | | Precentral_L |
| 6 | 21 | 6 | 10 | 5.52 | 1292 | Undefined |
| 6a | 21 | 15 | -2 | 5.21 | | Putamen_R |
| 7 | 42 | 12 | 26 | 5.27 | 861 | Frontal_Inf_Oper_R |
| 7a | 42 | 0 | 30 | 5.21 | | Undefined |

### 7. Correlations between the success metric and questionnaires

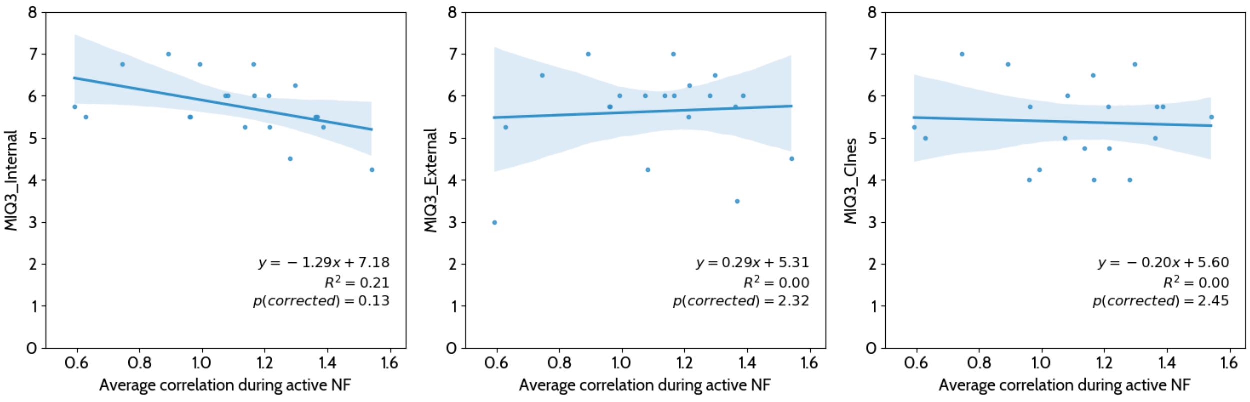


*Figure S5 - Correlating the MI-Q3 Internal, External, and Kinesthetic scores with the average correlation during active NF.*

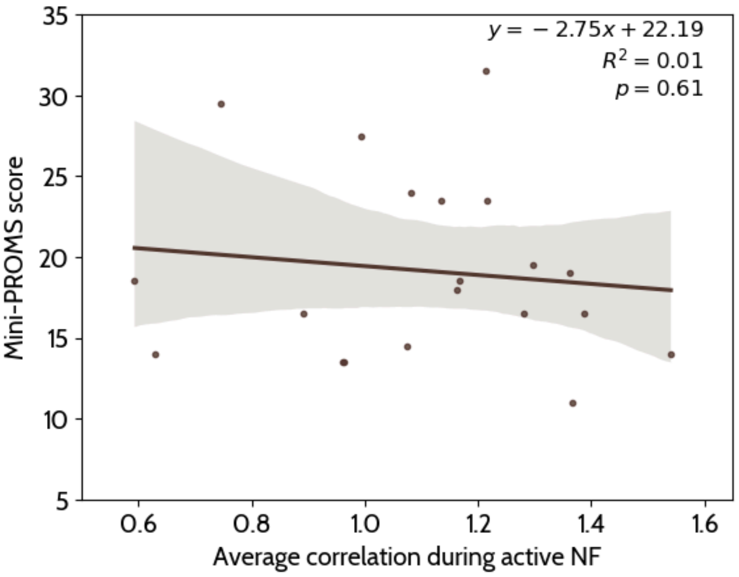


*Figure S6 - Correlating the Mini-PROMS score with the average correlation during active NF.*

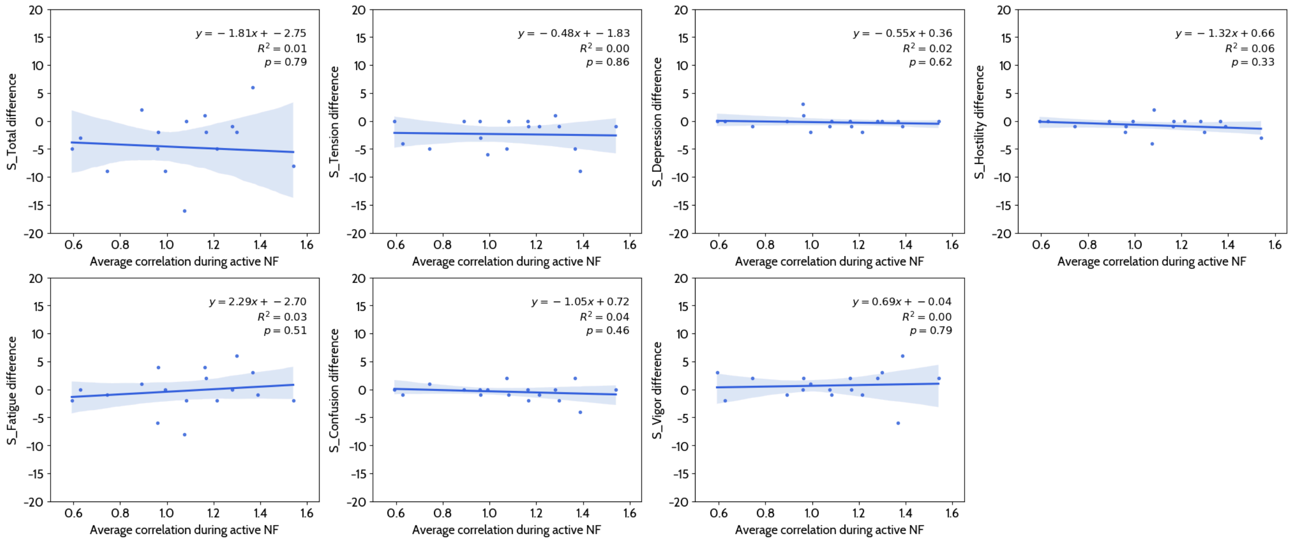


*Figure S7 - Correlating the POMS subscales difference between after and before the NF session with the average correlation during active NF.*

### 8. fMRIPrep methods

Results included in this manuscript come from preprocessing performed using fMRIPrep 24.0.1 (Esteban et al. (2019); Esteban et al. (2018); RRID:SCR_016216), which is based on Nipype 1.8.6 (K. Gorgolewski et al. (2011); K. J. Gorgolewski et al. (2018); RRID:SCR_002502).

#### 8.1. Preprocessing of B0 inhomogeneity mappings

A total of 1 fieldmaps were found available within the input BIDS structure for this particular subject. A B0 nonuniformity map (or fieldmap) was estimated from the phase-drift map(s) measure with two consecutive GRE (gradient-recalled echo) acquisitions. The corresponding phase-map(s) were phase-unwrapped with prelude (FSL None).

#### 8.2. Anatomical data preprocessing

A total of 1 T1-weighted (T1w) images were found within the input BIDS dataset. The T1w image was corrected for intensity non-uniformity (INU) with N4BiasFieldCorrection (Tustison et al. 2010), distributed with ANTs 2.5.1 (Avants et al. 2008, RRID:SCR_004757), and used as T1w-reference throughout the workflow. The T1w-reference was then skull-stripped with a Nipype implementation of the antsBrainExtraction.sh workflow (from ANTs), using OASIS30ANTs as target template. Brain tissue segmentation of cerebrospinal fluid (CSF), white-matter (WM) and gray-matter (GM) was performed on the brain-extracted T1w using fast (FSL (version unknown), RRID:SCR_002823, Zhang, Brady, and Smith 2001). Brain surfaces were reconstructed using recon-all (FreeSurfer 7.3.2, RRID:SCR_001847, Dale, Fischl, and Sereno 1999), and the brain mask estimated previously was refined with a custom variation of the method to reconcile ANTs-derived and FreeSurfer-derived segmentations of the cortical gray-matter of Mindboggle (RRID:SCR_002438, Klein et al. 2017). A T2-weighted image was used to improve pial surface refinement. Brain surfaces were reconstructed using recon-all (FreeSurfer 7.3.2, RRID:SCR_001847, Dale, Fischl, and Sereno 1999), and the brain mask estimated previously was refined with a custom variation of the method to reconcile ANTs-derived and FreeSurfer-derived segmentations of the cortical gray-matter of Mindboggle (RRID:SCR_002438, Klein et al. 2017). Volume-based spatial normalization to one standard space (MNI152NLin2009cAsym) was performed through nonlinear registration with antsRegistration (ANTs 2.5.1), using brain-extracted versions of both T1w reference and the T1w template. The following template was were selected for spatial normalization and accessed with TemplateFlow (24.2.0, Ciric et al. 2022): ICBM 152 Nonlinear Asymmetrical template version 2009c [Fonov et al. (2009), RRID:SCR_008796; TemplateFlow ID: MNI152NLin2009cAsym].

#### 8.3. Functional data preprocessing

For each of the 5 BOLD runs found per subject (across all tasks and sessions), the following preprocessing was performed. First, a reference volume was generated, using a custom methodology of fMRIPrep, for use in head motion correction. Head-motion parameters with respect to the BOLD reference (transformation matrices, and six corresponding rotation and translation parameters) are estimated before any spatiotemporal filtering using mcflirt (FSL , Jenkinson et al. 2002). The BOLD reference was then co-registered to the T1w reference using bbregister (FreeSurfer) which implements boundary-based registration (Greve and Fischl 2009). Co-registration was configured with six degrees of freedom. The aligned T2w image was used for initial co-registration.Several confounding time-series were calculated based on the preprocessed BOLD: framewise displacement (FD), DVARS and three region-wise global signals. FD was computed using two formulations following Power (absolute sum of relative motions, Power et al. (2014)) and Jenkinson (relative root mean square displacement between affines, Jenkinson et al. (2002)). FD and DVARS are calculated for each functional run, both using their implementations in Nipype (following the definitions by Power et al. 2014). The three global signals are extracted within the CSF, the WM, and the whole-brain masks. Additionally, a set of physiological regressors were extracted to allow for component-based noise correction (CompCor, Behzadi et al. 2007). Principal components are estimated after high-pass filtering the preprocessed BOLD time-series (using a discrete cosine filter with 128s cut-off) for the two CompCor variants: temporal (tCompCor) and anatomical (aCompCor). tCompCor components are then calculated from the top 2% variable voxels within the brain mask. For aCompCor, three probabilistic masks (CSF, WM and combined CSF+WM) are generated in anatomical space. The implementation differs from that of Behzadi et al. in that instead of eroding the masks by 2 pixels on BOLD space, a mask of pixels that likely contain a volume fraction of GM is subtracted from the aCompCor masks. This mask is obtained by dilating a GM mask extracted from the FreeSurfer's aseg segmentation, and it ensures components are not extracted from voxels containing a minimal fraction of GM. Finally, these masks are resampled into BOLD space and binarized by thresholding at 0.99 (as in the original implementation). Components are also calculated separately within the WM and CSF masks. For each CompCor decomposition, the k components with the largest singular values are retained, such that the retained components' time series are sufficient to explain 50 percent of variance across the nuisance mask (CSF, WM, combined, or temporal). The remaining components are dropped from consideration. The head-motion estimates calculated in the correction step were also placed within the corresponding confounds file.

The confound time series derived from head motion estimates and global signals were expanded with the inclusion of temporal derivatives and quadratic terms for each (Satterthwaite et al. 2013). Frames that exceeded a threshold of 0.5 mm FD or 1.5 standardized DVARS were annotated as motion outliers. Additional nuisance timeseries are calculated by means of principal components analysis of the signal found within a thin band (crown) of voxels around the edge of the brain, as proposed by (Patriat, Reynolds, and Birn 2017). All resamplings can be performed with a single interpolation step by composing all the pertinent transformations (i.e. head-motion transform matrices, susceptibility distortion correction when available, and co-registrations to anatomical and output spaces). Gridded (volumetric) resamplings were performed using nitransforms, configured with cubic B-spline interpolation.

#### 8.4. Functional data preprocessing

For each of the 5 BOLD runs found per subject (across all tasks and sessions), the following preprocessing was performed. First, a reference volume was generated, using a custom methodology of fMRIPrep, for use in head motion correction. Head-motion parameters with respect to the BOLD reference (transformation matrices, and six corresponding rotation and translation parameters) are estimated before any spatiotemporal filtering using mcflirt (FSL , Jenkinson et al. 2002). The estimated fieldmap was then aligned with rigid-registration to the target EPI (echo-planar imaging) reference run. The field coefficients were mapped on to the reference EPI using the transform. The BOLD reference was then co-registered to the T1w reference using bbregister (FreeSurfer) which implements boundary-based registration (Greve and Fischl 2009). Co-registration was configured with six degrees of freedom. The aligned T2w image was used for initial co-registration.Several confounding time-series were calculated based on the preprocessed BOLD: framewise displacement (FD), DVARS and three region-wise global signals. FD was computed using two formulations following Power (absolute sum of relative motions, Power et al. (2014)) and Jenkinson (relative root mean square displacement between affines, Jenkinson et al. (2002)). FD and DVARS are calculated for each functional run, both using their implementations in Nipype (following the definitions by Power et al. 2014). The three global signals are extracted within the CSF, the WM, and the whole-brain masks. Additionally, a set of physiological regressors were extracted to allow for component-based noise correction (CompCor, Behzadi et al. 2007). Principal components are estimated after high-pass filtering the preprocessed BOLD time-series (using a discrete cosine filter with 128s cut-off) for the two CompCor variants: temporal (tCompCor) and anatomical (aCompCor). tCompCor components are then calculated from the top 2% variable voxels within the brain mask. For aCompCor, three probabilistic masks (CSF, WM and combined CSF+WM) are generated in anatomical space. The implementation differs from that of Behzadi et al. in that instead of eroding the masks by 2 pixels on BOLD space, a mask of pixels that likely contain a volume fraction of GM is subtracted from the aCompCor masks. This mask is obtained by dilating a GM mask extracted from the FreeSurfer's aseg segmentation, and it ensures components are not extracted from voxels containing a minimal fraction of GM. Finally, these masks are resampled into BOLD space and binarized by thresholding at 0.99 (as in the original implementation). Components are also calculated separately within the WM and CSF masks. For each CompCor decomposition, the k components with the largest singular values are retained, such that the retained components' time series are sufficient to explain 50 percent of variance across the nuisance mask (CSF, WM, combined, or temporal). The remaining components are dropped from consideration. The head-motion estimates calculated in the correction step were also placed within the corresponding confounds file. The confound time series derived from head motion estimates and global signals were expanded with the inclusion of temporal derivatives and quadratic terms for each (Satterthwaite et al. 2013). Frames that exceeded a threshold of 0.5 mm FD or 1.5 standardized DVARS were annotated as motion outliers. Additional nuisance timeseries are calculated by means of principal components analysis of the signal found within a thin band (crown) of voxels around the edge of the brain, as proposed by (Patriat, Reynolds, and Birn 2017). All resamplings can be performed with a single interpolation step by composing all the pertinent transformations (i.e. head-motion transform matrices, susceptibility distortion correction when available, and co-registrations to anatomical and output spaces). Gridded (volumetric) resamplings were performed using nitransforms, configured with cubic B-spline interpolation.

Many internal operations of fMRIPrep use Nilearn 0.10.4 (Abraham et al. 2014, RRID:SCR_001362), mostly within the functional processing workflow. For more details of the pipeline, see the section corresponding to workflows in fMRIPrep's documentation.

#### 8.5. Copyright Waiver

The above boilerplate text was automatically generated by fMRIPrep with the express intention that users should copy and paste this text into their manuscripts unchanged. It is released under the CC0 license.

#### 8.6. References

Abraham, Alexandre, Fabian Pedregosa, Michael Eickenberg, Philippe Gervais, Andreas Mueller, Jean Kossaifi, Alexandre Gramfort, Bertrand Thirion, and Gael Varoquaux. 2014. "Machine Learning for

Neuroimaging with Scikit-Learn." Frontiers in Neuroinformatics 8. https://doi.org/10.3389/fninf.2014.00014.

Avants, B. B., C. L. Epstein, M. Grossman, and J. C. Gee. 2008. "Symmetric Diffeomorphic Image Registration with Cross-Correlation: Evaluating Automated Labeling of Elderly and Neurodegenerative Brain." Medical Image Analysis 12 (1): 26–41. https://doi.org/10.1016/j.media.2007.06.004.

Behzadi, Yashar, Khaled Restom, Joy Liau, and Thomas T. Liu. 2007. "A Component Based Noise Correction Method (CompCor) for BOLD and Perfusion Based fMRI." NeuroImage 37 (1): 90–101. https://doi.org/10.1016/j.neuroimage.2007.04.042.

Ciric, R., William H. Thompson, R. Lorenz, M. Goncalves, E. MacNicol, C. J. Markiewicz, Y. O. Halchenko, et al. 2022. "TemplateFlow: FAIR-Sharing of Multi-Scale, Multi-Species Brain Models." Nature Methods 19: 1568–71. https://doi.org/10.1038/s41592-022-01681-2.

Dale, Anders M., Bruce Fischl, and Martin I. Sereno. 1999. "Cortical Surface-Based Analysis: I. Segmentation and Surface Reconstruction." NeuroImage 9 (2): 179–94. https://doi.org/10.1006/nimg.1998.0395.

Esteban, Oscar, Ross Blair, Christopher J. Markiewicz, Shoshana L. Berleant, Craig Moodie, Feilong Ma, Ayse Ilkay Isik, et al. 2018. "fMRIPrep 24.0.1." Software. https://doi.org/10.5281/zenodo.852659.

Esteban, Oscar, Christopher Markiewicz, Ross W Blair, Craig Moodie, Ayse Ilkay Isik, Asier Erramuzpe Aliaga, James Kent, et al. 2019. "fMRIPrep: A Robust Preprocessing Pipeline for Functional MRI." Nature Methods 16: 111–16. https://doi.org/10.1038/s41592-018-0235-4.

Fonov, VS, AC Evans, RC McKinstry, CR Almli, and DL Collins. 2009. "Unbiased Nonlinear Average Age-Appropriate Brain Templates from Birth to Adulthood." NeuroImage 47, Supplement 1: S102. https://doi.org/10.1016/S1053-8119(09)70884-5.

Gorgolewski, K., C. D. Burns, C. Madison, D. Clark, Y. O. Halchenko, M. L. Waskom, and S. Ghosh. 2011. "Nipype: A Flexible, Lightweight and Extensible Neuroimaging Data Processing Framework in Python." Frontiers in Neuroinformatics 5: 13. https://doi.org/10.3389/fninf.2011.00013.

Gorgolewski, Krzysztof J., Oscar Esteban, Christopher J. Markiewicz, Erik Ziegler, David Gage Ellis, Michael Philipp Notter, Dorota Jarecka, et al. 2018. "Nipype." Software. https://doi.org/10.5281/zenodo.596855.

Greve, Douglas N, and Bruce Fischl. 2009. "Accurate and Robust Brain Image Alignment Using Boundary-Based Registration." NeuroImage 48 (1): 63–72. https://doi.org/10.1016/j.neuroimage.2009.06.060.

Jenkinson, Mark, Peter Bannister, Michael Brady, and Stephen Smith. 2002. "Improved Optimization for the Robust and Accurate Linear Registration and Motion Correction of Brain Images." NeuroImage 17 (2): 825–41. https://doi.org/10.1006/nimg.2002.1132.

Klein, Arno, Satrajit S. Ghosh, Forrest S. Bao, Joachim Giard, Yrjö Häme, Eliezer Stavsky, Noah Lee, et al. 2017. "Mindboggling Morphometry of Human Brains." PLOS Computational Biology 13 (2): e1005350. https://doi.org/10.1371/journal.pcbi.1005350.

Patriat, Rémi, Richard C. Reynolds, and Rasmus M. Birn. 2017. "An Improved Model of Motion-Related Signal Changes in fMRI." NeuroImage 144, Part A (January): 74–82. https://doi.org/10.1016/j.neuroimage.2016.08.051.

Power, Jonathan D., Anish Mitra, Timothy O. Laumann, Abraham Z. Snyder, Bradley L. Schlaggar, and Steven E. Petersen. 2014. "Methods to Detect, Characterize, and Remove Motion Artifact in Resting State fMRI." NeuroImage 84 (Supplement C): 320–41. https://doi.org/10.1016/j.neuroimage.2013.08.048.

Satterthwaite, Theodore D., Mark A. Elliott, Raphael T. Gerraty, Kosha Ruparel, James Loughead, Monica E. Calkins, Simon B. Eickhoff, et al. 2013. "An improved framework for confound regression and filtering for control of motion artifact in the preprocessing of resting-state functional connectivity data." NeuroImage 64 (1): 240–56. https://doi.org/10.1016/j.neuroimage.2012.08.052.

Tustison, N. J., B. B. Avants, P. A. Cook, Y. Zheng, A. Egan, P. A. Yushkevich, and J. C. Gee. 2010. "N4ITK: Improved N3 Bias Correction." IEEE Transactions on Medical Imaging 29 (6): 1310–20. https://doi.org/10.1109/TMI.2010.2046908.

Zhang, Y., M. Brady, and S. Smith. 2001. "Segmentation of Brain MR Images Through a Hidden Markov Random Field Model and the Expectation-Maximization Algorithm." IEEE Transactions on Medical Imaging 20 (1): 45–57. https://doi.org/10.1109/42.906424.
