## Supplementary material for "A novel music-based real-time fMRI neurofeedback interface modulates interhemispheric connectivity and enhances mood": CRED-NF Checklist

### CRED-nf checklist summary

*06 May, 2025*

**Manuscript title:** Music as a real-time fMRI neurofeedback interface for modulating interhemispheric connectivity: effects on mood and recruitment of the putamen and insula

**Corresponding Author:** Alexandre Sayal

| Item No. | Checklist item | Manuscript Details |
|---|---|---|
| **Pre-experiment** | | |
| 1a | Pre-register experimental protocol and planned analyses | This study was preregistered on OSF (https://doi.org/10.17605/OSF.IO/AHXNB). |
| 1b | Justify sample size | A sample of 22 healthy participants (13 females, mean age 33 ± 6 years, range 24-42 years) was recruited for this study. Based on previous research on neurofeedback, we aimed for an effect size of 1.09 ( =0.05) for modulation activity difference between active and sham conditions. Based on G*Power, the estimated sample size per group was 19 participants. |
| **Control groups** | | |
| 2a | Employ control group(s) or control condition(s) | A crossover design was implemented: half of the participants received sham feedback during the first two runs, while the other half received sham feedback during the last two runs. The feedback shown during the sham NF runs was randomly generated. |
| 2b | When leveraging experimental designs where a double-blind is possible, use a double-blind | *NA: A double-blind was not appropriate for this experiment* |
| 2c | Blind those who rate the outcomes | *Those who rated the outcome were not blind to group assignment* |
| | Blind those who analyse the data | *Those who analysed the data were not blind to group assignment* |
| 2d | Examine to what extent participants and experimenters remain blinded | After the MRI session, participants were asked to describe their experiences, including their perceived performance and feedback contingency during the runs. They were also asked to identify the best and worst runs regarding these parameters. Finally, the runs in which they received sham feedback were disclosed. |
| 2e | In clinical efficacy studies, employ a standard-of-care intervention group as a benchmark for improvement | *NA: This is not a clinical efficacy study* |
| **Control measures** | | |

| | | |
|---|---|---|
| 3a | Collect data on psychosocial factors | After recruitment and before the MRI acquisition session, participants completed a series of questionnaires. The first questionnaire assessed each participant's compatibility with the MRI environment to ensure safety and compliance. Next, we characterized the sample based on three parameters: musical training, handedness, and motor imagery ability. Participants completed the Mini-PROMS questionnaire (Zentner and Strauss 2017), which assessed musical ability, and provided the number of years of formal music training. To assess handedness, we used a questionnaire (Cohen 2008) based on (Oldfield 1971). To evaluate motor imagery ability, participants completed the MIQ-3 questionnaire (Mendes et al. 2016; Williams et al. 2012), which screens for visual and kinesthetic imagery capabilities. |
| 3b | Report whether participants were provided with a strategy | Participants then watched a video explaining the neurofeedback experiment. This video detailed the number and type of sequences, the paradigm of the two tasks, and included a guided opportunity to practice motor imagery while listening to feedback music. (...) The participants were asked to imagine tapping the fingers of their hands during the 'Motor imagery' condition and to attentively listen to the music during the 'Music' condition. |
| 3c | Report the strategies participants used | *The strategies participants used were not recorded or not reported in the manuscript* |
| 3d | Report methods used for online-data processing and artifact correction | Real-time analysis was performed in Turbo-BrainVoyager v4. After the localizer run, the ROIs in the bilateral premotor cortices of each participant were defined based on the activation map of the contrast between 'Motor imagery' and 'Rest' conditions. The statistical threshold was set to t-value $> 3.1$ ($p < 0.001$ uncorrected). Anatomical landmarks of the precentral gyrus, visible when superimposing the anatomical T1w image, were also considered in this definition. For all functional runs, the preprocessing steps included motion correction, inter-run alignment, and detrending. An iterative General Linear Model (GLM) was used to estimate the activation maps for each run. The design matrices included predictors for each of the experimental conditions and the six motion parameters (translation and rotation in the X, Y, and Z axes) as confound predictors. |
| 3e | Report condition and group effects for artifacts | *Condition and group effects for artifacts were not measured, or not reported in the manuscript* |
| **Feedback specifications** | | |
| 4a | Report how the online-feature extraction was defined | The 'Motor imagery' was accompanied by musical feedback, which was calculated using the windowed correlation (8-second window) between the activity of the left and right premotor cortices. |

| | | |
|---|---|---|
| 5a | Report neurofeedback regulation success based on the feedback signal | The basis for the feedback provided, and also the modulation target for the participants, was Pearson's correlation between the BOLD signals of the left and right PMCs, as defined in the localizer run. These correlations were estimated in an 8-second sliding window across time during the acquisition. In Figure 8, we display the average of the estimated correlations across subjects for the active and sham runs during the NF 'Motor imagery' and 'Rest' blocks. |
| 5b | Plot within-session and between-session regulation blocks of feedback variable(s), as well as pre-to-post resting baselines or contrasts | *The manuscript does not plot within-session and between-session regulation blocks of feedback variable(s), as well as pre-to-post resting baselines or contrasts* |
| 5c | Statistically compare the experimental condition/group to the control condition(s)/group(s) (not only each group to baseline measures) | One of the questions of this feasibility study regards the activation patterns of contingent feedback vs. sham. For this comparison, we considered the thresholded group-level activation maps (p=0.05 with Bonferroni's correction, k>10) of active and sham feedback runs and defined a binary mask including the significant voxels for each of them. Then, we summed these masks to obtain a map of overlap between the two conditions - in practice, in each voxel, we show if the activity was significant for both conditions, only for active, or only for sham feedback. This map provides an overview of the pattern of network recruitment, while we look more closely at the reward and learning circuits. |
| **Outcome measures - behaviour** | | |
| 6a | Include measures of clinical or behavioural significance, defined a priori, and describe whether they were reached | The assessment of participants' mood before and after the MRI NF session was performed based on the self-reports of the POMS questionnaire. A significant decrease was found for the Tension sub-scale score (Mdn before = 3, Mdn after = 0, p = 0.012, W = 4) and for the Total score (Mdn before = 104, Mdn after = 98, p = 0.012, W = 28) - Figure 10. The results for the remaining sub-scales do not present a significant difference between time points and are presented in Figure S4. |
| 6b | Run correlational analyses between regulation success and behavioural outcomes | Here, we assessed changes in mood as an outcome measure, via the POMS self-reporting questionnaire. We found that the overall score and the Tension subscale decreased after the NF session, which are signs of mood improvement. This scale has been found to change after NF in previous studies reporting a full NF intervention in autism spectrum disorder using a visual interface exploiting facial expressions (Direito et al. 2021) and in major depressive disorder using a visual thermometer (Mehler et al. 2021; 2018). In our case, none of the differences are correlated with the success metric (Supplementary Figure S7). (...) Regarding the Mini-PROMS and MIQ-3 questionnaires, we found no statistically significant correlations with the success metric (Figures S5 and S6). |
| **Data storage** | | |

| | | |
|---|---|---|
| 7a | Upload all materials, analysis scripts, code, and raw data used for analyses, as well as final values, to an open access data repository, when feasible | All code used in this study is available in the GitHub repository (https://github.com/CIBIT-UC/musicnf-novelinterface). The dataset, formatted in BIDS, can be accessed at Zenodo (https://doi.org/10.5281/zenodo.14803374). |
